# In silico saturation mutagenesis of cancer genes

**DOI:** 10.1101/2020.06.03.130211

**Authors:** Ferran Muiños, Francisco Martinez-Jimenez, Oriol Pich, Abel Gonzalez-Perez, Nuria Lopez-Bigas

## Abstract

Extensive bioinformatics analysis of datasets of tumor somatic mutations data have revealed the presence of some 500-600 cancer driver genes. The identification of all potential driver mutations affecting cancer genes is essential to implement precision cancer medicine and to understand the interplay of mutation probability and selection in tumor development. Here, we present an in silico saturation mutagenesis approach to identify all driver mutations in 568 cancer genes across 66 tumor types. For most cancer genes the mutation probability across tissues --underpinned by active mutational processes-- influences which driver variants have been observed, although this differs significantly between tumor suppressor and oncogenes. The role of selection is apparent in some of the latter, the observed and unobserved driver mutations of which are equally likely to occur. The number of potential driver mutations in a cancer gene roughly determines how many mutations are available for detection across newly sequenced tumors.

## Introduction

Tumors follow a darwinian evolution through the interplay between somatic variation and selection (Greaves and Maley, 2012; McGranahan and Swanton, 2017; Stratton, 2011; Stratton et al., 2009). Through the sequencing of tens of thousands of tumor whole exomes over one and a half decade of cancer genomics we have come to understand the contribution of different mutational processes and chromatin features to the emergence of coding variation (Alexandrov et al., 2013, 2020; Frigola et al., 2017; Gonzalez-Perez et al., 2019; Lawrence et al., 2013; Nik-Zainal et al., 2012; Pich et al., 2018; Polak et al., 2014, 2015; Sabarinathan et al., 2016; Stamatoyannopoulos et al., 2009; Supek and Lehner, 2019). Researchers have also been successful at tracking the occurrence of selection at the level of genes through the detection of signals of positive selection (Arnedo-Pac et al.; Dees et al., 2012; Dietlein et al., 2020; Gonzalez-Perez and Lopez-Bigas, 2012; Kamburov et al., 2015; Lawrence et al., 2013; Martínez-Jiménez et al., 2019; Niu et al., 2016; Tamborero et al., 2013a; Vogelstein et al., 2013). Between 500 and 600 cancer genes, which likely constitute an important fraction of the compendium of all genes capable of driving cancer through point mutations have thus been identified (Bailey et al., 2018; Gonzalez-Perez et al., 2013; Rubio-Perez et al., 2015; Sondka et al., 2018; Tamborero et al., 2013b). In tumors sequenced we have observed a fraction of all possible point mutations in these genes, some of which, but not all, are known to be capable of driving tumorigenesis (Martincorena et al., 2017; Sabarinathan et al., 2017). In general, our knowledge on how selection acts at the level of individual mutations in cancer genes across different tissues is still limited.

To be able to study selection at the level of individual mutations a novel approach is required that classifies all possible mutations in a gene, independently of their probability of occurrence, as drivers or passengers. Furthermore, it would be desirable that such classification yields human-readable results, which help researchers point at the key features defining driver mutations in a cancer gene. Instead of relying on functional impact metrics (Kircher et al., 2014; Pollard et al., 2010), this method should measure the ability of a mutation to drive tumorigenesis. Moreover, unlike existing methods to identify individual driver mutations (Tokheim and Karchin, 2019), this approach should take into account that, in different genes, and across tissues, different features are relevant to define selection on individual mutations.

This problem has been addressed before through experiments of saturation mutagenesis, in which all possible mutants of a cancer gene are generated and their impact on protein function (Kakudo et al., 2005; Kato et al., 2003; Kawaguchi et al., 2005) or cell viability (Findlay et al., 2018; Mighell et al., 2018) are assessed. These experiments possess obvious technical and economic hurdles. Furthermore, due to limitations imposed by the experimental setup, these approaches do not directly measure the tumorigenic potential of mutations, but rather some proxy, such as their functional impact. For instance, in certain tumor suppressor genes, saturation mutagenesis experiments have been conducted in haploid human cells to identify mutations that abrogate cell viability (Findlay et al., 2018). Only scattered mutagenesis assays have been carried out that actually assess the tumorigenic potential of mutations affecting cancer genes, restricted to few cell types, which do not represent the wide spectrum of tissue-specific constraints. Generalizing them to cover hundreds of cancer genes across cell types representing different tissues would be a herculean task.

With the aim of circumventing these inconveniences, we have designed a gene-tissue specific in silico saturation mutagenesis approach. To that end, we leverage the mutational features identified across 568 cancer genes in 28,000 samples of 66 tumor types to build an ensemble of gene and tumor type-specific machine learning models that capture the making of their driver mutations. Models are trained on observed and synthetically generated mutations across genes and cohorts of tumors. With this ensemble of models we identified the potential driver mutations across all possible variants of each cancer gene, and repeated this process for all tumor types in which the gene was identified as driver. Since the predictions of the models are interpretable, we are able to learn what features describe a driver mutation in a cancer gene in a specific tissue. Using the distribution of all potential driver mutations in each cancer gene, the portion they represent of all possible mutations and the fraction of them that we observe in each cancer gene, we can explore the interplay between the probability of occurrence of mutations and selective constraints across tissues. We thus address fundamental questions of cancer genomics, such as how much the profile of potential driver mutations of a cancer gene differ between tumor types as a reflection of the selective constraints acting in different tissues. We also explore the reason why potential driver mutations in cancer genes remain unobserved in different tumor types, and we assess the likelihood to observe them through sequencing new tumors.

## Results

### An approach to in silico saturation mutagenesis of cancer genes

In silico saturation mutagenesis of a cancer gene in a tissue consists in classifying each possible nucleotide change along its sequence, whether it has been observed or not in sequenced tumors, as driver or passenger (Fig. 1a). To this end we created boostDM, a machine learning approach that effectively exploits the patterns of mutational features computed in 568 cancer genes across 28,000 samples of 66 tumor types (Table S1; www.intogen.org) to build models that, learning the properties that define driver mutations of a cancer gene in each tumor type, are able to classify all the mutations they can possibly bear into drivers and passengers (Fig. 1b; Supp. Note). Mutational features include the significant clustering of mutations along the primary structure of the protein or on its three-dimensional structure (Arnedo-Pac et al.; Tokheim et al., 2016), the significant enrichment for mutations in functional domains (Martínez-Jiménez et al., 2019), the unexpected number of observed mutations of different consequence types (Martincorena et al., 2017), but also some specific to each mutation, such as the site conservation (Pollard et al., 2010) or whether it is post-translationally modified (PTM) (Bateman et al., 2017).

**Figure 1.**
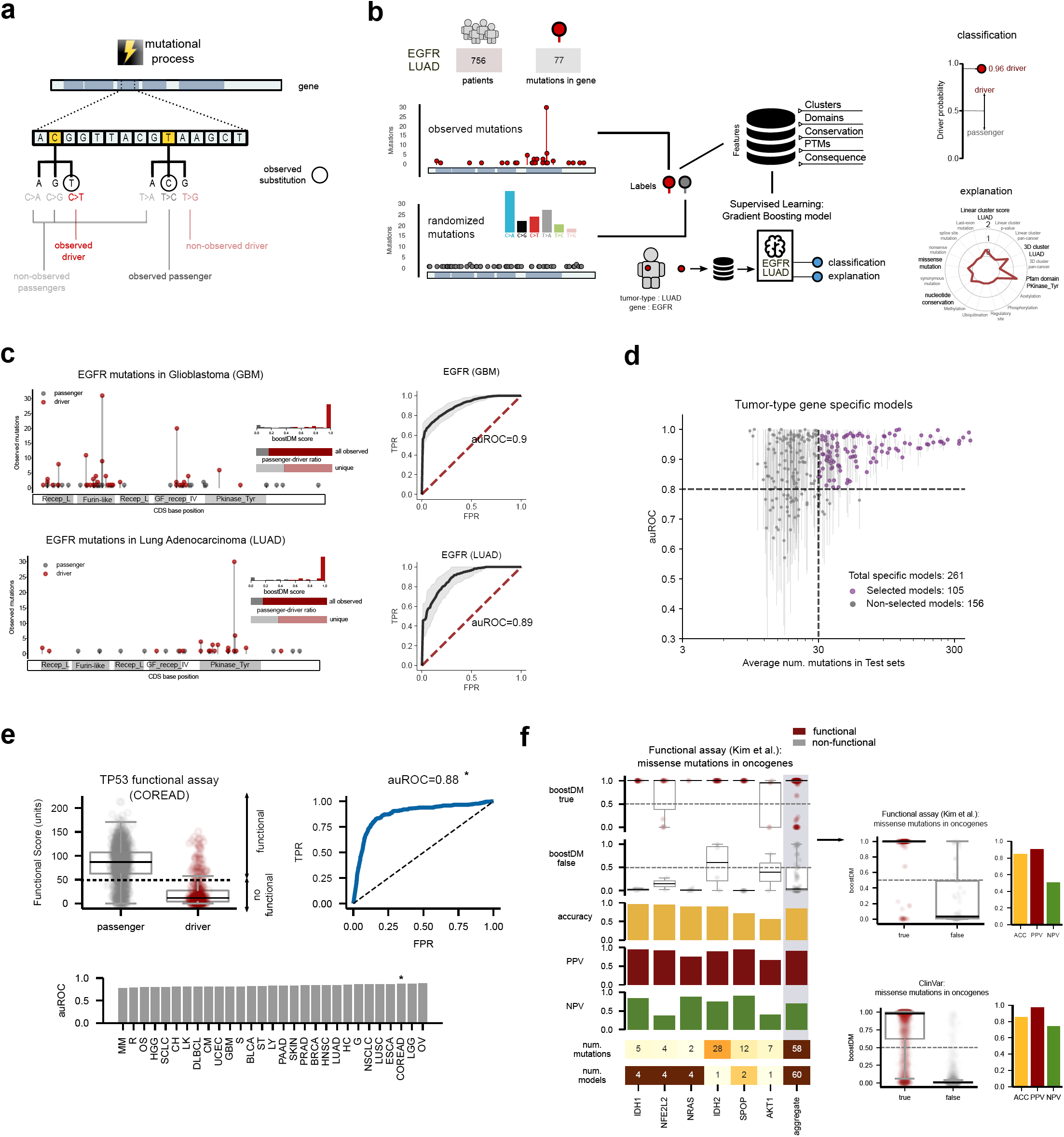
A novel approach for in silico saturation mutagenesis of cancer genes. a) Conceptual representation of in silico saturation mutagenesis in a cancer gene. Every possible mutation in the gene may be classified as a potential driver (either observed or unobserved) or a potential passenger. b) Schematic representation of the construction of a cancer gene-tumor type specific drivers identification model (EGFR across a cohort of lung adenocarcinomas) by the boostDM method. Among 756 patients in the cohort, 133 EGFR mutations have been identified. These constitute the positive set to train the model. The negative set (randomized mutations) is integrated by 6,650 EGFR possible mutations sampled following the mutational tri-nucleotide frequencies observed across the cohort. All observed and randomized mutations are annotated with pre-computed gene-level (ref IntOGen) and mutation-level features. The resulting Gradient Boosting model can then be used to score the 133 observed mutations (driver probability) and classify them into drivers and passengers. An explanation of the combined contribution of mutational features to the classification of each mutation is also obtained. PTM: post translational modification. c) Classification of observed mutations in EGFR across glioblastomas and lung adenocarcinomas. Inset panels show the distribution of boostDM driver score, fraction of all mutations, and fraction of unique mutations classified as drivers across EGFR in each tissue. Cross-validation ROC curves for each model are obtained by different subsets of observed mutations and re-generated randomized mutations. The black line represents the mean ROC, and in grey we represent the area spanned by the ROC curves across 50 classifiers trained with different train-test splits from the same data. d) Distribution of auROC (y-axis) of 313 cancer gene-tumor type specific models. The x-axis values represent the number of observed mutations used to train the model. The dot in each distribution represents the median auROC value for the model, with the line comprising its 95% confidence intervals. e) The in silico saturation mutagenesis approach applied to TP53 yields results that are highly coherent with those obtained through experimental saturation mutagenesis. Clockwise from top left plot: i) distribution of the functional score of all TP53 mutations classified as drivers (red) or passengers (gray) by the colorectal specific model; ii) ROC curve derived from this classification; iii) auROC resulting from the classification of mutations using different tumor type specific mutations. f) Performance of different cancer gene-tumor type specific models on the classification of validated functional and non-functional mutations. The left panel represents the performance of pooled tumor type specific models for each cancer gene with at least 1 functional and 1 neutral mutations experimentally validated by (Kim et al., 2016). The two top columns exhibit the boostDM driver score of functional (red) and non-functional (gray) mutations in cancer genes. The rightmost column represents the performance of models on aggregated mutations. A summary of the performance across all missense mutations in oncogenes is represented in the top right plot. The bottom right plot contains a summary of the performance of specific models on missense somatic pathogenic and germline neutral variants in oncogenes manually annotated from the literature (Landrum et al., 2018).

Building one machine learning model representing the features of driver mutations across all driver genes would fail to capture differences in tumorigenic mechanisms among them. The differences in the mutational patterns of archetypal tumor suppressor genes and oncogenes constitute a salient example of this. The former exhibit abnormal rates of truncating mutations or inactivating missense mutations distributed along the entire protein sequence, while the latter frequently accumulate mutations only in specific domains, which in some cases cluster in only one or two amino acid residues (Davoli et al., 2013). Moreover, differences in tumorigenic mechanisms reflecting upon diverse mutational features are probably the norm amongst cancer genes. We thus reasoned that constructing models that capture the mutational features of individual cancer cancer gene-tumor types pairs, rather than a single global model is necessary to carry out the in silico saturation mutagenesis assay.

To build a cancer gene-tumor type specific model two sets of bona fide i) tumorigenic (positives) and ii) passenger (negatives) mutations are required. Representative datasets with sufficient numbers of experimentally validated driver and passenger are inexistent. Nevertheless, it is possible to accurately compute the fraction of observed mutations that are in excess in a cancer gene across tumors in a cohort, given their expected number accounting for the frequency of tri-nucleotide substitutions and other determinants of the background mutation rate (Martincorena et al., 2017; Sabarinathan et al., 2017). These excess mutations, unexplained by the background mutation rate of the cancer gene are drivers. Thus, in cancer genes with a very high fraction of excess mutations, say above 0.85, most observed mutations across tumors are drivers, and constitute a good positive training set. Using all observed mutations as a positive training set has the added advantage of avoiding potential selection biases that may be present in sets of validated oncogenic variants. For example, 77 observed mutations in EGFR across 756 lung adenocarcinomas (LUAD) constitute the positive set to train a EGFR-LUAD specific model (Fig. 1b).

Passenger mutations occur along the sequence of a cancer gene following the probability of each site to mutate. Therefore, we reasoned that a set of synthetic mutations drawn from the gene following the frequency of tri-nucleotide changes observed in the cohort would constitute a good negative training set. Each mutation in these two sets is annotated with a set of features (Fig. 1b). (Fig. 1b, left; Supp. Note.). A model (gradient boosting machine) is then trained using these positive and negative sets of annotated mutations for EGFR-LUAD (Supp. Note.). In all, 261 cancer gene-tumor type combinations fulfilled the criterion of an excess of at least 85% of observed over expected non silent mutations, that allowed us to build specific models for them. For each particular observed mutation, the model yields a score (ranging from 0 to 1), which may be interpreted as a probability that it is a driver. Mutations with scores above 0.5 are labeled as drivers, while the rest are considered passengers (Fig. 1b, top right). BoostDM also produces an assessment of the contribution of all mutational features to this classification (Fig. 1b, bottom right).

The outcome and performance of EGFR models specific for two different tumor types are illustrated in Figure 1c. Of all EGFR-affecting mutations observed across glioblastomas (GBM) and lung adenocarcinomas (LUAD), boostDM identifies 137 and 65 mutations as drivers, respectively. Fifty partial views of the set of driver and synthetic mutations were selected to train the models and the remaining mutations to test them in a 70/30 cross-validation (Supp. Note). Their performance can be measured as the area under the Receiver Operating Characteristic (ROC) curve (auROC). Their values (0.9 in the case of the GBM model and 0.89 for the LUAD model) reflect that these two gene-tumor type specific boostDM models classify observed mutation as drivers or passengers with high accuracy.

The performance of specific models increases with the number of mutations employed to train them. We regard as good quality specific models those with test sets of at least 30 mutations and yielding auROC above 0.8 (Fig. 1d). As more datasets of mutations across tumor cohorts become available, we foresee that the number of high-quality specific models will subsequently increase. Overall, 105 models specific for a cancer gene and a tumor type fulfill this condition. To classify the mutations in cancer gene-tumor type combinations whose specific models do not fulfill the conditions set above, more general models can be built by pooling the mutations of related malignancies (according to the oncotree hierarchy; Table S2; Supp. Note; Fig. S1a-f) into meta-cohorts (Zehir et al., 2017). Moreover, general models representing the features of mutations across all tumor suppressors or oncogenes (or all cancer genes) in cohorts or meta-cohorts can also be built. The most suitable non-specific model to classify the mutations of a gene in a tumor type is decided on the basis of the accuracy of models of decreasing specificity (Supp. Note.). With a combination of specific and non-specific models we are then able to identify driver mutations affecting 568 cancer genes across 64 tumor types, obtaining an average of 2.1 drivers per tumor (1.1-3.1) across non-hypermutated samples (see Methods). This is close (Fig. S2a,b) to a recent estimation of the number of driver mutations per tumor across a smaller set of malignancies based on the quota of positive selection (Martincorena et al., 2017).

An independent assessment of the performance of boostDM may be carried out through the classification by several TP53-tumor type specific models of variants generated in the course of an experimental saturation mutagenesis assay of this gene. In this experiment, the functional impact of all TP53 non-silent mutations on its ability to transactivate four promoter constructs in yeast was measured (Kakudo et al., 2005; Kato et al., 2003; Kawaguchi et al., 2005). The separation of these mutations into drivers and passengers shows great agreement with their experimentally measured functional impact, with all tested models yielding auROC above 0.75 (Fig. 1e). A similar comparison can be carried out with another saturation mutagenesis experiment (Supp. Note.)

Saturation mutagenesis experiments are cost and labor intensive, and thus very rare. Therefore, we also assessed the performance of gene-tumor type specific models in the task of distinguishing mutations in 22 cancer genes whose functionality has been experimentally verified, collected from two independent sources (KIM and BERGER) (Berger et al., 2016; Kim et al., 2016). We classified the missense mutations affecting oncogenes employing all gene-tumor type specific models. The classifications yielded by all models were pooled and the accuracy, positive predictive value (PPV) and negative predictive value (NPV) computed (Fig. 1f; Fig. S2c). The same analysis was done for mutations affecting tumor suppressor genes (Fig. S2d). For most cancer genes in both datasets, the models showed high accuracy. The global accuracy (polling the classification by different specific models of mutations in all genes) reaches 0.84 (mutations in oncogenes in KIM dataset), 0.77 (mutations in oncogenes in BERGER dataset) and 0.80 (mutations in tumor suppressors in KIM dataset). Moreover, we verified that models accurately distinguish pathogenic (or likely pathogenic) somatic mutations from benign (or likely benign) germline variants in cancer genes manually collected and curated by the ClinVar resource (Landrum et al., 2018) across 80 cancer genes, respectively (Fig. 1f; Fig. S2e).

Driver mutations affecting different cancer genes are best described by the dissimilar contribution of mutational features to their classification by a specific model. For example, the residence in a mutational cluster detected on the three-dimensional structure (Tokheim et al., 2016) of EGFR makes a very important contribution to the classification of the A289D mutation as driver by the GBM specific model (Fig. 2a). Smaller contributions are made by the location of this mutation in a linear mutational cluster (Arnedo-Pac et al.), the conservation of the site across orthologs (Pollard et al., 2010), its occurrence in the Furin-like domain (Finn et al., 2010) of EGFR and the fact that it is a missense mutation (McLaren et al., 2010) in a protein with an excess of missense mutations (Martincorena et al., 2017). The classification of driver mutations affecting other cancer genes across cancer types by their respective specific models is based on the combined importance of mutational features (Fig. 2b). As expected, the tructating nature of mutations affecting certain tumor suppressor genes proves decisive to their classification as drivers (APC R213* and RB1 E51* and a splice-affecting mutation of TP53). However, different features, such as their location within a cluster defined on the three-dimensional structure of the protein (TP53 K132R), their function as a phosphorylation site (VHL S80R) or its occurrence within a key functional domain (CIC R202W) play a key role in the identification of other tumor-suppressor affecting driver mutations. Being part of a mutational cluster, defined either linearly or in three dimensions provides an important contribution to the classification of exemplary driver mutations affecting two oncogenes (PIK3CA E542K and CTNNB1 S45P).

**Figure 2.**
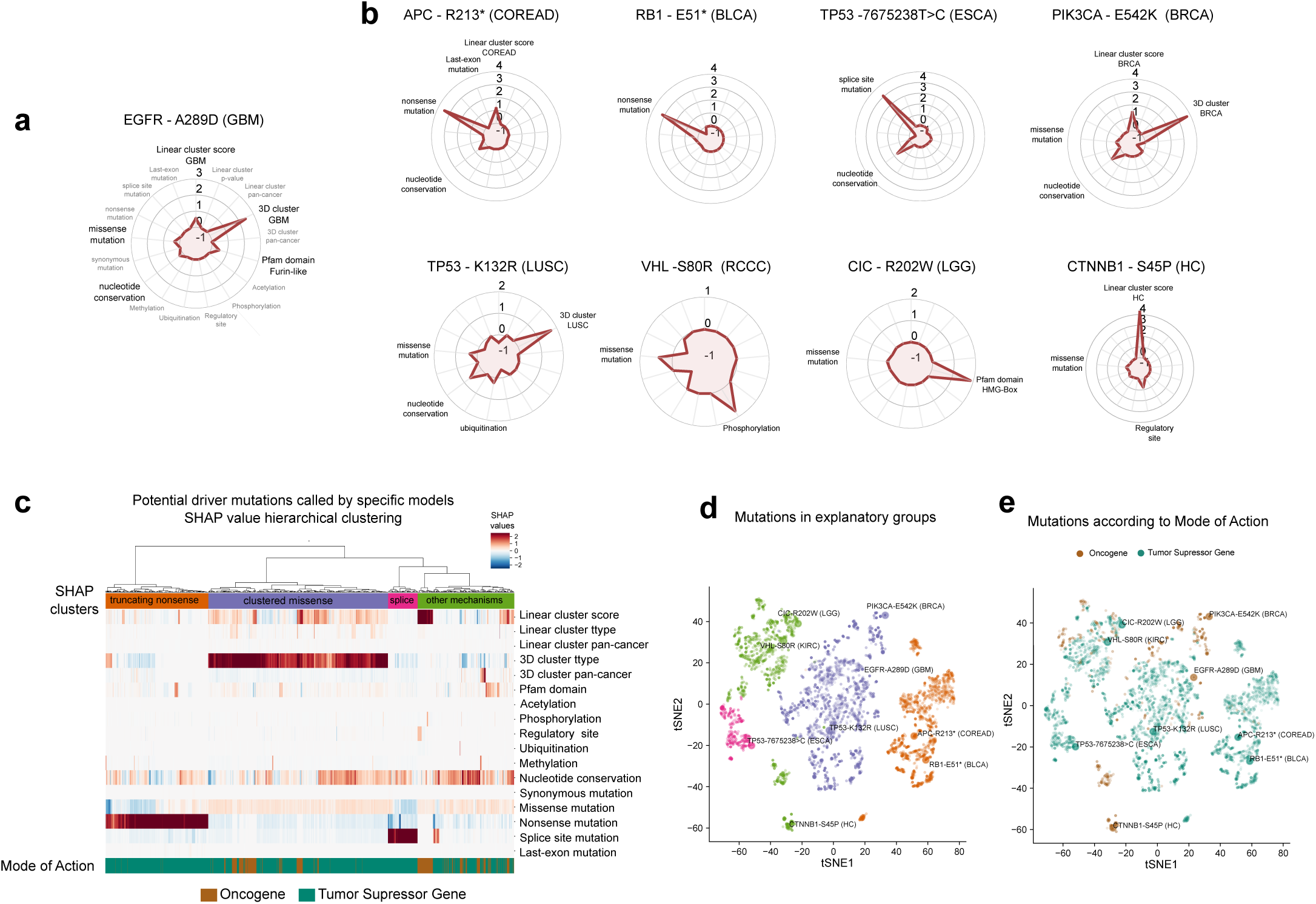
The making of driver mutations. a) Radial plot representation of the combined contribution of mutational features to the classification of the EGFR A289D as a driver in glioblastomas. Mutational features are represented around the most external concentric circle, and the red line tracks the relative contribution of each to the classification. Radial values are SHAP values in logit scale, i.e., positive (resp. negative) values push forecast above (resp. below) 0.5 probability that the mutation is a driver. Mutational features with significant positive contribution to the classification of this mutation as driver appear in black font; those with null or negative contribution, appear in gray font. b) Combined contribution of mutational features to the classification of mutations of different genes among several tumor types as drivers. c) Relative contribution of all mutational features to the classification of all driver mutations by the 148 specific models. Mutations are grouped through a hierarchical clustering (colors and dendrogram above the heatmap). d,e) Clustering of driver mutations according to the combined contribution of mutational features to their classification in a t-SNE two-dimensional projection. Mutations are colored according to (d) their grouping in c) or (e) the mode of action of the cancer genes they affect.

Importantly, it is ultimately a complex interaction between different features within the specific model, rather than a single feature that determines the identification of a driver mutation. These combinations for all mutations identified as drivers by specific models are laid out in Figure 2c. Certain groups of mutations, the classification of which is driven by particular combinations of features, become apparent (Fig. 2c,d). These groups correspond to different contributions of features to the identification of driver mutations affecting tumor suppressor and oncogenes, but also different classes within each of these two major groups (Fig. 2e).

In summary, gene-tumor type specific models trained by boostDM are capable of accurately identifying driver mutations in cancer genes. The outcome of the models can also be used to understand which combination of mutational features defines each driver mutation.

### Heterogeneous landscape of observed and unobserved driver mutations across cancer genes and tumor types

We then carried out an in silico saturation mutagenesis (evaluating every possible nucleotide change) of 78 cancer genes across 40 malignancies employing gene-tumor type specific models, thus obtaining the profile of potential driver mutations in a cancer gene in a tissue. Figure 3 (a-d) and Figure S3 (a,b) illustrate this profile for six cancer genes-tumor type combinations.

**Figure 3.**
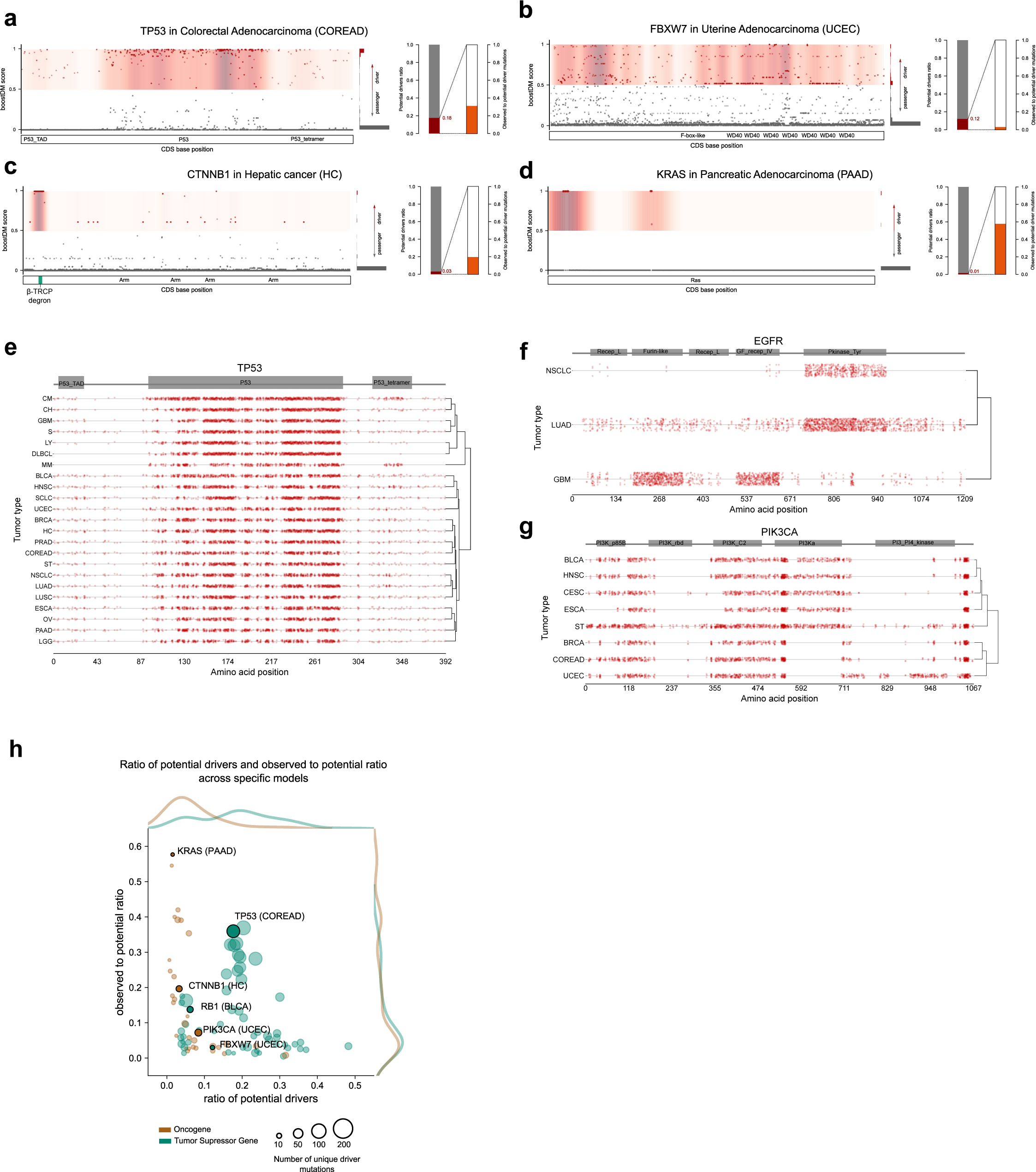
In silico saturation mutagenesis of cancer genes. a-d) Result of applying the in silico saturation mutagenesis approach to four cancer genes. All possible mutations in each cancer gene (dots) are represented in their relative position of the gene coding sequence (x-axis). Relevant functional protein domains are represented within each gene body. Potential driver mutations appear in red and potential passenger mutations, gray. The concentration of driver mutations at different regions of the protein is represented as a density. The distribution of boostDM driver scores of all possible mutations appears at the right side of each saturation mutagenesis plot. Two bar plots illustrate the potential drivers ratio (red on gray) and the ratio of observed-to-potential driver mutations (orange on gray). e-g) Profile of potential driver mutations of three cancer genes across several tumor types. h) Two-dimensional plot representing the relationship between the ratio of potential drivers (y-axis) and the observed-to-potential ratio (x-axis). Cancer gene-tumor type combinations are represented as circles colored according to their mode of action and with their size proportional to the number of unique driver mutations identified by the in silico saturation mutagenesis approach. The distribution of the two ratios for tumor suppressor and oncogenes is shown along each corresponding axis.

The distribution of potential driver mutations along the sequence of the gene is starkly different between the three tumor suppressors and the three oncogenes shown in Figure 3 (a-d) and Figure S3 (a,b). Potential driver mutations of KRAS across pancreatic adenocarcinomas cluster around the two known mutational hotspots in codons 12-13 and 61 (refs). In the case of CTNNB1 in hepatocellular carcinomas, most potential driver mutations are clustered within and around the degron of the protein (refs). Several clusters, including those at well-known 542-545 and 1047 codons group most potential driver mutations identified in PIK3CA across uterine adenocarcinomas. In the three tumor suppressors, potential driver mutations appear more widespread along the gene sequence, although regions with clusters of driver mutations --some of them overlapping domains, as in the case of TP53 across colorectal adenocarcinomas-- can still be appreciated.

A first question that we can address with the results of the in silico saturation mutagenesis is what fraction of all possible mutations in a cancer gene are potentially drivers. In the six exemplary cancer genes, this ratio of potential drivers (red-gray bar by each gene saturation mutagenesis plot) falls between 1% (KRAS in pancreatic adenocarcinoma) and 18% (TP53 in colorectal adenocarcinoma). That is, only 1% of all possible nucleotide changes in KRAS are identified as potential drivers by boostDM, because their feature combinations resemble those encoded in the model specific of pancreatic adenocarcinomas. The in silico saturation mutagenesis of the three tumor suppressors (Fig. 3a-c) reveals higher ratios of potential drivers (6%, 12% and 18%) than those of the three oncogenes (Fig. 3d-f; Fig. S3c; 1%, 3% and 7%). This trend is observed for all tumor suppressors and oncogenes assessed, with the ratio of potential drivers ranging from less than 1% to 40%. It is not surprising that oncogenes exhibit a smaller ratio of potential drivers than tumor suppressor genes since intuitively, more sites are available for loss-of-function than for gain-of-function. The profiles of potential driver mutations of a cancer gene differ between tumor types (Fig. 3e-g and Fig. S3d,e). The origin of these differences may be the different selective constraints specific to each tissue that shape the landscape of fitness-gain positions in the sequence of a cancer gene.

We next asked what fraction of all potential driver mutations of a cancer gene in a tissue we have observed in sequenced tumors. We refer to this quantity as the observed-to-potential ratio and is represented by the orange fraction of the second bar by each gene saturation mutagenesis plot. Strikingly, in the exemplary cancer genes in the Figures (with the exception of KRAS in pancreatic adenocarcinomas), the observed-to-potential ratio is below 0.5. This means that across cohorts of these malignancies, we have observed fewer than half of all potential driver mutations identified through the in silico saturation mutagenesis in these cancer genes. This fraction ranges between 0.5% and 57% across all cancer genes (Fig. S3f). Despite their smaller ratio of potential drivers, oncogenes do not show a trend towards higher observed-to-potential ratios (Fig. S3g). Although a few oncogenes (as KRAS in pancreatic adenocarcinomas) do exhibit remarkably high observed-to-potential ratio (above any tumor suppressor) no clear differences between tumor suppressors and oncogenes are apparent. Furthermore, no clear relationship is appreciable between the ratio of potential drivers and the observed-to-potential ratio of cancer genes (Fig. 3g). This is probably because while the former quantity is determined solely by the availability of gain-of-function or loss-of-function sites, the second is influenced both by the strength of selection on sites and the probability of individual driver mutations.

### The influence of mutation probability on observed driver mutations

We set out to disentangle the interplay between the probability of individual driver mutations and their fitness gain of observed driver mutations in cancer genes. To that end, we computed the probability of occurrence of all potential driver mutations (mutation probability; Supp. Data) in a cancer gene in a tumor type obtained via its in silico saturation mutagenesis. This mutation probability is based on the tri-nucleotide frequency profile of all mutations observed in the cohort (Supp. Methods). For each driver gene in a tissue we then define two metrics of the probability bias between observed and unobserved driver mutations (Fig. 4a). The first corresponds to the area under a ROC curve (auROC) tracking the separation of observed and unobserved driver mutations based on mutation probability. The second is the fold-change (logFC) of the median mutation probability of observed and unobserved mutations. If the mutation probability plays an important role in defining which driver mutations we have observed, we expect a positive mutation probability bias for cancer genes, that is, values of auROC above 0.5 and positive mutation probability logFC values.

**Figure 4.**
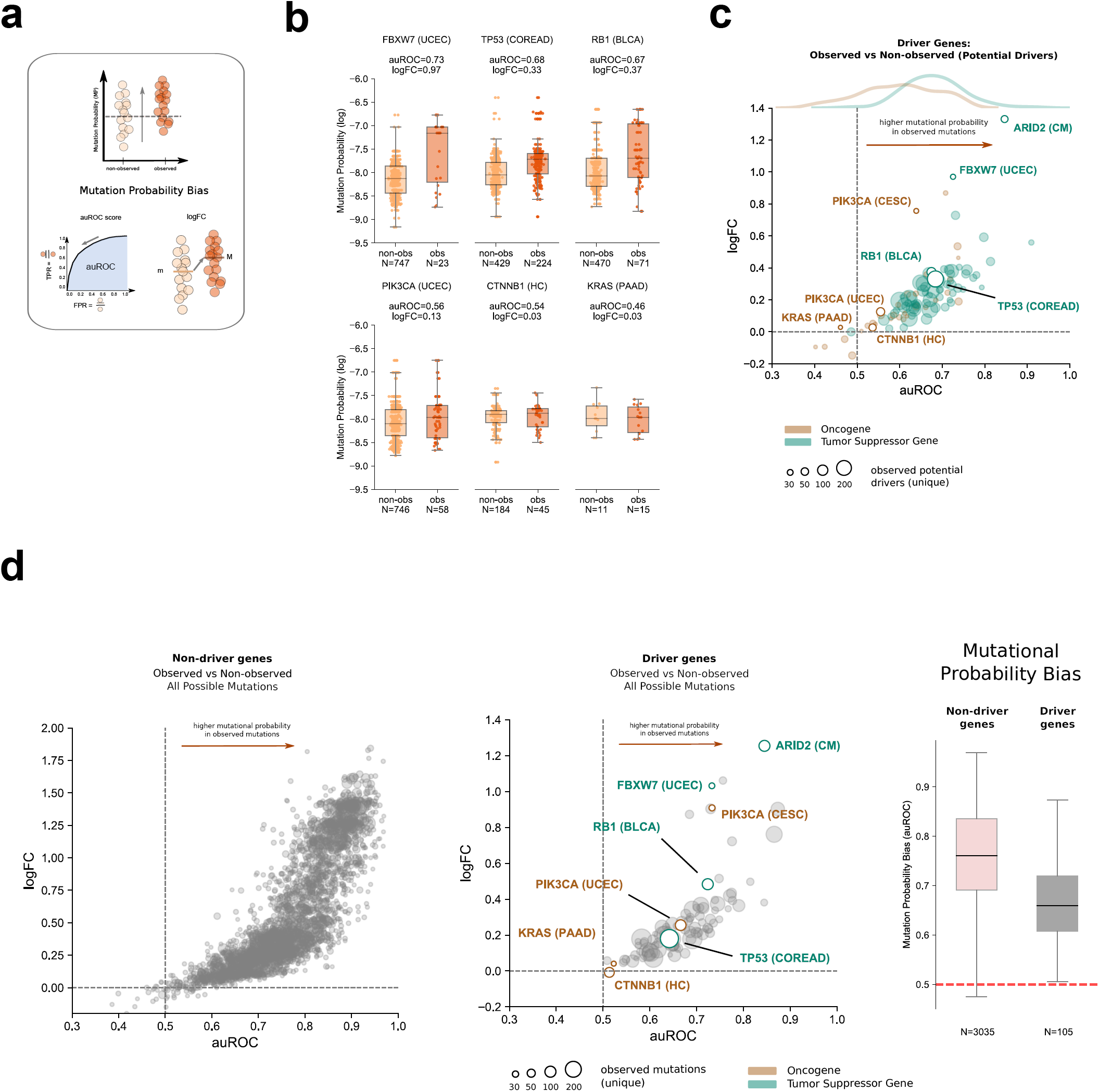
The mutation probability bias of cancer genes. a) Conceptual representation of the mutation probability bias of a cancer gene and its calculation. Top: the probability of occurrence of observed (dark orange) and unobserved mutations (light orange) is computed (mutation probability); bottom left: scanning the values of mutation probability, a ROC curve may be constructed, the area under which (auROC) assesses the efficiency of the separation of observed and unobserved mutations; bottom right: a fold change (logarithm, or logFC) of the median mutation probabilities of both distributions may also be used to assess the efficiency of their separation. b) Mutational probability bias (auROC and logFC) of six exemplary cancer genes. The distribution of mutation probability of observed and unobserved driver mutations are represented as boxplots with the points representing the actual mutations upon them. c) Two-dimensional plot representing the mutation probability bias of 105 cancer gene-tumor type combinations with specific models. The x-axis represents the auROC and the y-axis, the logFC. Cancer genes are represented as circles colored according to their mode of action and their size following their number of unique observed driver mutations. d) The first graph presents the mutation probability bias of 3,035 non-cancer gene-tumor type combinations (with at least 10 mutations reported in intOGen). The mutation probability bias is computed with all possible mutations (canonical transcript) of each gene. The second graph presents the mutation probability bias of 105 cancer gene-tumor type combinations, computed with all possible mutations of each cancer gene. In the rightmost graph, the distributions of mutation probability bias (auROC) presented in the two previous graphs are compared.

Among the six exemplary genes of previous figures, we found that the three tumor suppressors have a strong mutation probability bias, with their auROC values between 0.67 and 0.73 and logFC between 0.31 and 0.95 (Fig. 4b). On the other hand, this is much weaker for the three oncogenes. PIK3CA across uterine adenocarcinomas and CTNNB1 in hepatocellular carcinomas display almost no mutation bias at all with auROC values of 0.55 and 0.54, and logFC very close to zero. A slight inverse mutational probability bias is observed for driver mutations of KRAS in pancreatic adenocarcinomas (auROC=0.46 and logFC close to zero). Generalizing this analysis across cancer genes we observed that most exhibit a strong mutation probability bias (Fig. 4c). Most exceptions to this behavior --weak or non-existent bias-- correspond to oncogenes (Fig. 4c; Fig. S4a,b). This means that across most cancer genes the probability of occurrence of driver mutations has at least some influence on which of them are observed.

To gain a better understanding of the role of mutational probability in shaping the observed-to-potential ratio, we next computed the mutational bias across all possible mutations in non cancer genes (Methods; Fig. 4d, first panel). As expected, in this scenario in which selection is not expected to play a preponderant role, the mutational bias is much stronger than that of all mutations in cancer genes is comparatively smaller (Fig. 4d, second and third panels). This difference in the mutation probability bias between non-cancer and cancer genes is likely due to the contribution of the selective advantage provided by driver mutations to cells.

In summary the action of positive selection on specific mutations in cancer genes and the probability on which mutations are observed in tumors shape the landscape of observed mutations. The observed absence of mutation probability bias in some oncogenes could thus be explained by an extreme effect of selection on relatively few available gain-of-function mutations, which are not among the most frequently mutated.

### Mutational signatures define driver mutations available in different tissues

The probability of occurrence of mutations is determined by the mutational processes active in a tissue. Therefore, we next looked at their relationship with the likelihood of observation of certain driver mutations across tumor types (Table S3; Methods).

We first focused on two close mutational hotspots of PIK3CA in breast tumors (affecting aminoacid residues 542 and 545; Fig. 5a). For all possible mutations affecting nine codons of PIK3CA around these hotspots, we computed the contribution of three mutational processes (identified by mutational signatures 1, 2 and 5) to the mutational probability in average breast tumors (see Methods and Supp. Note). The two most frequent mutations in these hotspots (E542K and E545K) have higher probability to occur than neighboring mutations (top panel), due to the activity of the APOBEC-driven mutational process (middle panel). A third possible mutation (affecting codon 543) of PIK3CA, also within the APOBEC context conferring high probability yields a synonymous change and is thus not observed in tumors.

**Figure 5.**
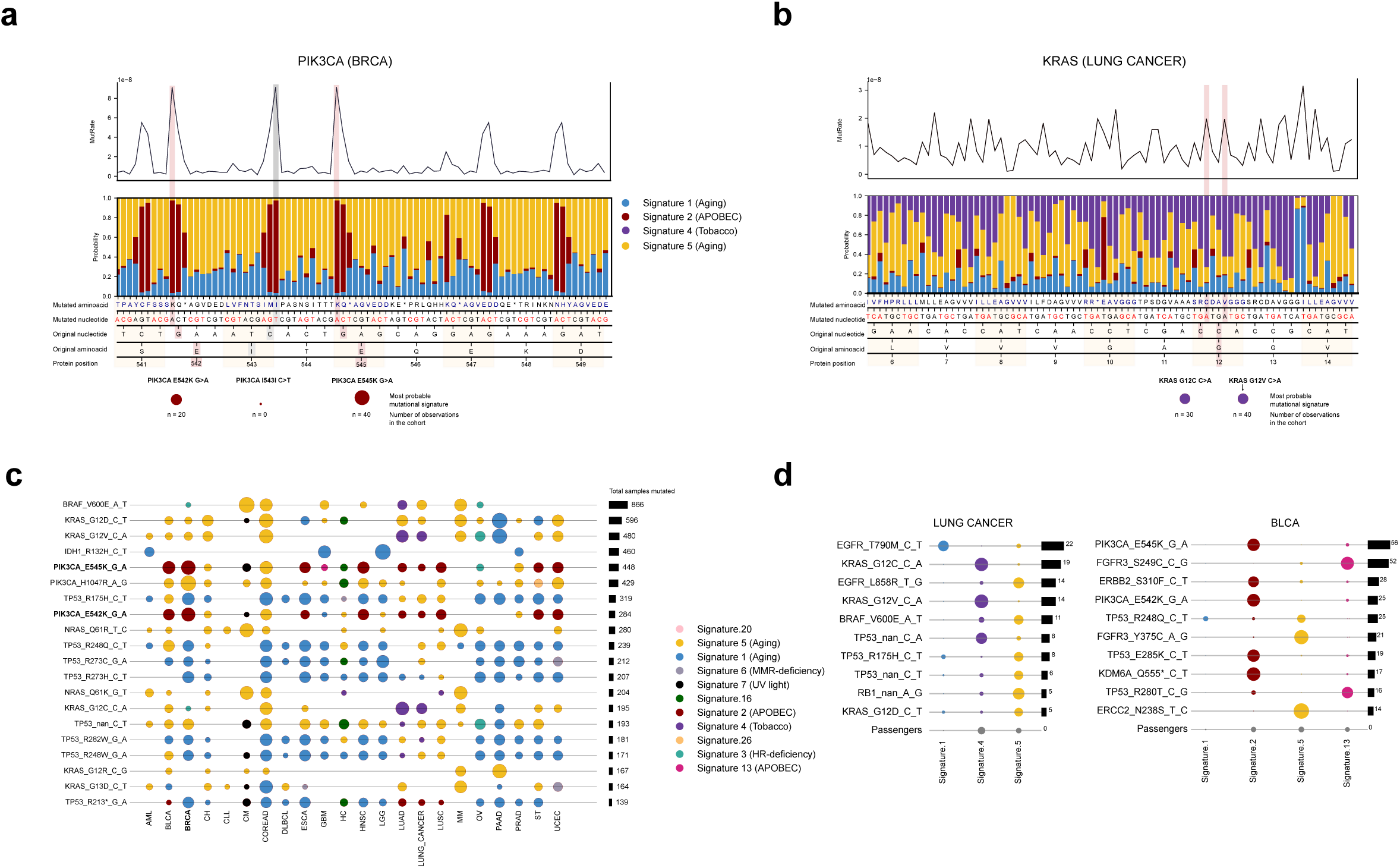
Mutational processes and driver mutations across tissues. a) Probability of all possible mutations in a nine amino acid long segment of the PIK3CA sequence in breast adenocarcinomas. The top plot represents the probability of all possible nucleotide changes (MutRate, see Methods) in this segment of the protein sequence. In the bottom plot, the relative contribution of three mutational processes active across breast tumors to the probability of each possible mutation is represented. Each possible mutation is labelled below the graph with the reference and alternative nucleotide and amino acid. b) Same as a) for all possible mutations in a nine amino acid long segment of the KRAS sequence in lung tumors. c) Most likely mutational processes contributing several frequent driver mutations across tissues. The contribution of 11 mutational signatures to the occurrence of a driver mutation in each cancer type is represented as a circle colored according to the signature and with its size proportional to the frequency of the driver mutation in the malignancy. Bars at the right of the plot track the total frequency of each driver mutation. d) Relative contribution of mutational processes active across lung tumors or breast adenocarcinomas to driver mutations frequent in these malignancies. The baseline contribution of each mutational process to the mutation burden of tumors of each tissue (labelled passengers) is represented at the bottom row of each plot.

A different landscape of mutational processes activity surrounds the hotspot in codon 12 of KRAS in lung tumors (including changes of the wild-type glycine to cysteine or valine; Fig. 5b). Many neighboring positions (with tobacco or aging signatures contexts) possess mutational probability values similar to those of this codon. Nevertheless, although they show similar occurrence probability, they provide no increase in fitness to lung cells and are thus virtually never observed in tumors.

Driver mutations that appear frequently across different tumor types are either contributed by mutational processes common to many tissues or contributed by different mutational processes active across them (Fig. 5c). For example, while the NRAS Q61R mutation across tumor types represented in the Figure appears to be contributed by the ubiquitous aging-related signature 5, the aforementioned PIK3CA recurrent mutations in codons 542 and 545 (and a third hotspot in position 1047) are contributed by either the APOBEC related signature 2, the aging related signature, the UV-light signature or signature 16 depending on the cancer type.

The interaction between accessibility of a nucleotide change in a given context and the activity of different processes determines their contribution to the landscape of driver mutations in a tissue (Fig. 5d). For example, in lung tumors, the well-known driver mutations KRAS G12V and G12C are most likely contributed by the tobacco related signature, highly active in this tissue in smokers. On the other hand, the gatekeeping EGFR T790M is contributed with high probability by the aging related signature 1, despite its relatively lower activity (represented by the probability of contributing passenger mutations in the tissue), due to its context preference. In another example, in bladder adenocarcinomas, despite comparable activity of both APOBEC related signatures, their different context preference determines that certain driver mutations are most likely contributed by one or the other.

In summary, the mutational processes active in a tissue determine which driver mutations are available, and this may constitute a factor limiting the occurrence of driver mutations across cancer types.

### Observing new driver mutations in freshly sequenced tumors

The fact that some potential driver mutations have never been observed in cancer may be a consequence of the limited number of tumor whole exomes sequenced to date. Low observed-to-potential ratios across many cancer genes across tumor types suggest there is still a large room for the discovery of new driver mutations through sequencing. Therefore, it seems reasonable to assume that as more tumors of different cancer types are probed for somatic mutations, the observed-to-potential ratio of driver mutations of cancer genes shall increase. But, exactly how likely is it that we observe such an increase in the future for different cancer genes?

To answer this question, we carried out a systematic random subsampling of the tumors of different cancer types and fitted the expected number of unique driver mutations of each gene observed within the resulting random subsets of increasing sample size (see Methods). Then, a mutational discovery index was computed, ranging from 0 to 1, following the slope of the fitting curve at the point that corresponds to the actual number of samples of each cohort. This mutational discovery index tracks the unlikeliness of observing new unique mutations affecting the gene by sequencing more tumor samples.

Their values for the six exemplary cancer genes introduced in previous figures appear in Figure 6a-f. The three oncogenes exhibit larger mutational discovery index values (0.79-0.98) than the three tumor suppressor genes (0.3-0.7). For CTNNB1 in hepatocellular carcinomas and KRAS in pancreatic adenocarcinomas, the discovery of unobserved driver mutations has already plateaued for tumor cohorts with the sample size currently available in IntOGen. In other words, the number of observable driver mutations of CTNNB1 is approaching discovery saturation. On the other hand, with larger tumor cohorts, currently unobserved driver mutations of RB1, FBXW7 and TP53 are expected to continue appearing. Overall, oncogenes possess significantly higher mutational discovery index values than tumor suppressor genes (Fig. S5a).

**Figure 6.**
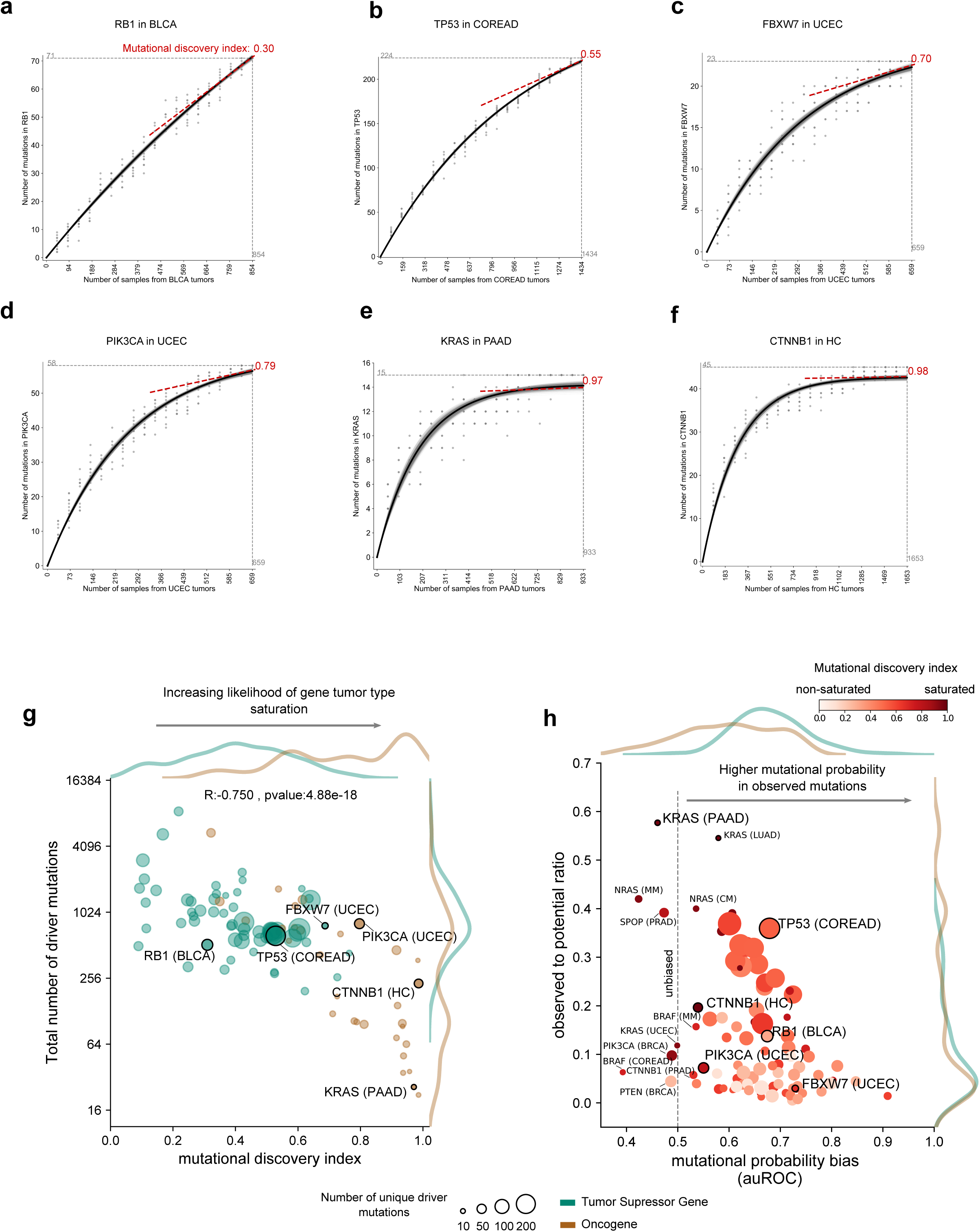
Potential for the discovery of new driver mutations across cancer genes. a-f) Mutational discovery index of six exemplary cancer genes. The dots in each plot represent the number of unique driver mutations identified across random subsamples drawn from each cohort, with the black curve representing the best exponential fit to them. A tangent to the best fit curve at the x-axis value corresponding to the actual sample size of the cohort is represented as a broken red line. The mutational discovery index (see Methods) derived from the slope of this tangent line is shown at each plot. g) Two-dimensional plot representing the relationship between the total number of potential driver mutations (y-axis) and the mutational discovery index (x-axis) of 110 cancer genes with specific models. Each cancer gene is represented as a circle colored by its mode of action and with size proportional to the number of unique observed driver mutations. The distribution of both values for tumor suppressor and oncogenes is shown along the axes. The pearson correlation coefficient representing the relationship between the two quantities and its p-value are included in the graph. h) Two-dimensional plot representing the relationship between the observed-to-potential ratio (y-axis) and the mutation probability bias (x-axis) of cancer genes with 105 specific models. Each cancer gene is represented as a circle colored following its mutational discovery index and with size proportional to the number of unique observed driver mutations. The distribution of both values for tumor suppressor and oncogenes is shown along the axes.

The mutational discovery index among cancer genes negatively correlates with the number of potential driver mutations they bear (Fig. 6g). Cancer genes with 100 or fewer potential driver mutations --all of them, oncogenes-- have reached or are very close to reaching discovery saturation with currently available tumor cohorts. As the number of potential driver mutations increases into the hundreds or thousands, the probability to discover one that has not been observed yet increases. This relationship, which holds for all observed mutations in cancer genes (Fig. S5b), constitutes further validation of the in silico saturation mutagenesis approach.

Interestingly, cancer genes with weak or non-existent mutation probability bias tend to show higher mutational discovery index values, irrespective of their observed-to-potential driver mutations ratio (Fig. 6h). Take, for example, the cases of CTNNB1 and KRAS, with almost identical mutational discovery index (0.97, CTNNB1 and 0.98, KRAS), but with KRAS exhibiting almost four-fold higher observed-to-potential driver mutations ratio. Given the virtual absence of mutation probability bias, the explanation is not that unobserved mutations occur at significantly lower probabilities than the ones we have already observed. It is possible that, although all potential KRAS or CTNNB1 driver mutations identified by the in silico saturation mutagenesis approach possess the features that are common to tumorigenic mutations, only a few of them confer the actual cell transforming capability. Alternatively, if all identified potential driver mutations are actually tumorigenic, some of them may confer a much stronger selective advantage than the rest, making the cells acquiring them more likely to initiate tumorigenesis. On the other hand, most tumor suppressors with medium to low mutational discovery index values (such as TP53 across colorectal adenocarcinomas) exhibit a strong mutation probability bias. It is thus reasonable to conclude that in their case, a number of unobserved driver mutations with lower probability of occurrence may still be detected through sequencing.

## Discussion

In a little over four decades of cancer genetics research since the discovery of the first cancer genes and driver mutations, the development of cancer genomics has brought about the possibility to uncover the complete compendium of cancer genes across tumor types (Bailey et al., 2018; Gonzalez-Perez et al., 2013; Rubio-Perez et al., 2015; Tamborero et al., 2013b). Despite sequencing tens of thousands of tumors over a span of roughly 15 years, only a small fraction of all possible mutations in cancer genes have been observed. Uncovering the landscape of all potential driver mutations in cancer genes across tissues is key to interpret the genomes of newly sequenced tumors in the clinical setting (Chakravarty et al., 2017; Griffith et al., 2017; Tamborero et al., 2018). It’s also essential to understand the interplay between mutation probability and selection in the profile of observed mutations in driver genes in tumors. Here, we have addressed these questions through a new in silico saturation mutagenesis approach. BoostDM, the method we developed to carry out this in silico saturation mutagenesis constitutes a first step towards this goal. One key innovation of this method is that specific models for cancer genes and tissues are trained, thus capturing the differences in the features that define driver mutations across genes and tumor types. BoostDM demonstrated high accuracy in distinguishing between validated driver and passenger mutations across cancer genes. Moreover it exhibited high accuracy when benchmarked against an experimental saturation mutagenesis assay and independent validated pathogenic and benign mutations across cancer genes. Importantly, models trained by boostDM are not black boxes. Instead, they produce easy-to-interpret interpretations of the rationale behind the classification of each mutation.

Nevertheless, there is ample room to improve its performance. The construction of boostDM models is based on the systematic collection of mutations in cancer genes across tumor types and the calculation of their mutational features that we carry out via the IntOGen pipeline (www.intogen.org). Therefore, as more datasets of tumor somatic mutations are released into the public domain and the calculation of new mutational features is incorporated into the IntOGen pipeline, higher-quality models of driver mutations across cancer genes will be obtained by boostDM. The expansion in the number of datasets of tumor somatic mutations will also provide more cancer gene-tumor type specific models, also contributing to increase the accuracy of classification of driver mutations. Furthermore, higher-quality models will also yield more complete and nuanced explanations of the features that define driver mutations across cancer genes and tissues. Importantly, since the driver score of a mutation does not take into account any information about the context of the tumor sample under analysis (such as other mutations or non-genetic features) it cannot be taken as a certainty of its tumorigenic role. Rather, it is to be interpreted as a measure of its similarity with mutational features extracted from the analysis of thousands of tumors.

The strong anticorrelation observed between the number of potential driver mutations and the mutational discovery index across cancer genes provides further support to the accuracy of the in silico saturation mutagenesis. Nevertheless, deviations from the trend may bear testimony to the aforementioned potential for refinement of the models. Consider the case of CTNNB1, used as illustration throughout Results. Most mutations affecting the B-TRCP degron in CTNNB1 (Martínez-Jiménez et al., 2019; Mészáros et al., 2017) are deemed drivers by the boostDM hepatocellular carcinoma specific model, driven primarily by their overlap of a cluster (Fig. S3c). Nevertheless, probably only some of them cause some (or, alternatively greater) perturbation to its recognition by the E3-ubiquitin ligase. It is precisely in this case that improvement of the models, through new features capturing subtler effects of mutations may refine the in silico saturation mutagenesis further.

We envision that the in silico saturation mutagenesis approach will become particularly relevant for the interpretation of newly sequenced tumor exomes (or panels) in the clinical or research settings (Tamborero et al., 2018). As the mutational discovery index indicates, for the vast majority of cancer genes, many yet unobserved mutations are potentially drivers. Therefore, counting with a reliable method to classify variants of unknown significance --in particular newly observed mutations-- is of paramount importance. To support this interpretation, we make the results of the in silico saturation mutagenesis available to researchers through the IntOGen platform (www.intogen.org/boostdm).

## Acknowledgements

We acknowledge important technical contributions to the development of the in silico saturation mutagenesis approach and to the manuscript by Loris Mularoni, Iker Reyes-Salazar, EIectra Tapanari and Claudia Arnedo-Pac. N.L-B. acknowledges funding from the European Research Council (consolidator grant 682398) and Spanish Ministry of Economy and Competitiveness (SAF2015-66084-R, MINECO/FEDER, UE). IRB Barcelona is a recipient of a Severo Ochoa Centre of Excellence Award from the Spanish Ministry of Economy and Competitiveness (MINECO; Government of Spain) and is supported by CERCA (Generalitat de Catalunya). O.P. is the recipient of a BIST PhD fellowship supported by the Secretariat for Universities and Research of the Ministry of Business and Knowledge of the Government of Catalonia, and the Barcelona Institute of Science and Technology (BIST). The results shown here are in whole or part based upon data generated by the TCGA Research Network, the Pan-Cancer Analysis of Whole Genomes (PCAWG), the cBioPortal, the Hartwig Medical Foundation, the International Cancer Genomes Consortium (ICGC), the St. Jude pediatric hospital, the Pediatric cBioPortal, TARGET projects, the BEAT AML study, and several other studies scattered throughout the scientific literature. This publication and the underlying study have been made possible partly on the basis of the data that Hartwig Medical Foundation has made available to the study.

## Author contributions

In silico saturation mutagenesis conceptualization: F.M., F.M-J., O.P., A.G-P. and N.L-B. Development of boostDM and mutation probability calculations: F.M. Downstream analyses and figure preparation: F.M., F.M-J., O.P. Analysis and discussion of results: F.M., F.M-J., O.P., A.G-P., N.L-B. Data collection: O.P., F.M-J., F.M. BoostDM pipeline development: F.M-J., F.M. Project supervision: A.G-P., N.L-B. Manuscript preparation: F.M., F.M-J., O.P., A.G-P., N.L-B.

## Methods

### Data Source: Cohorts of Sequenced Tumours

Somatic single nucleotide variants (SNV) from post-processed (i.e., after removal of hypermutated samples, multiple samples from the same donor, etc.) intOGen cohorts (release 1 February 2020) were used as a dataset of input observed mutations.

The release encompasses 28,076 samples with 203,003,747 somatic mutations from 221 cohorts of 66 different cancer types. For further information about the cohorts please refer to https://www.intogen.org/.

### In silico saturation mutagenesis

To conduct an in-silico saturation mutagenesis type of analysis, we developed a systematic learning approach (boostDM) intended to annotate and explain point mutations in cancer driver genes likely involved in tumorigenesis. This section briefly sketches the boostDM method. For a comprehensive account, please refer to the Supplementary Note in Supplemental Information.

BoostDM delineates a supervised learning strategy based on observed mutations in sequenced tumours and their site-by-site annotation with mutational features, comparing observed mutations in genes for which the observed-to-expected ratio is high enough with randomly selected mutations following the trinucleotide mutational probability, in terms of a reduced collection of mutational features. The method essentially looks into the protein coding sequence of the genome as all mutations considered map to the canonical transcripts in protein coding genes according to the Ensembl Variant Effect Predictor (VEP.92) (McLaren et al., 2010).

For each gene and tumor-type context, the method assigns a model on the basis of a model selection strategy based on cross-validation. Each such model is a collection of expert classifiers that reach a consensus probability score with an aggregator that intends to correct for the systematic bias of under-confident classifiers. The classifiers used are boosted tree models trained with a logistic binary objective loss function on subsets of the data. Furthermore, the tree-like structure of the models allow additive explanation models to be built, by additively splitting the forecast for each individual mutation in terms of the relative contributions (using Shapley Additive Explanations or SHAP values) of the features used. Thus the models can learn from training sets of annotated mutations, yield predictions for observed or unobserved annotated mutations and provide an explanation model in the form of average SHAP values (Lundberg and Lee, 2017) for the prediction at each individual mutation.

### Number of drivers per sample

Estimates of the average number of driver mutations per sample among the observed per sample were given using two different approaches: dNdScv (Martincorena et al., 2017) and boostDM (Supp. Fig. S2a).

### dNdScv

To compute the number of mutations in excess (i.e., the difference between the number of mutations observed and the number of mutations expected according to a neutral selection model) in cancer driver genes across the analyzed cohorts, we resorted to the methodology laid out by dNdScv (Martincorena et al., 2017).

Briefly, dNdScv provides a gene-specific and estimation of the ratio of non-synonymous to synonymous substitutions (dN/dS) that is corrected by i) chromatin features explaining regional variability of neutral mutation rate, ii) the consequence type of the substitutions and iii) the mutational processes operative in the tumor. For the purpose of our analysis we computed the excess of non-synonymous substitutions by aggregating the excess of missense, nonsense and splicing-affecting mutations.

Upon the estimation w of the dN/dS for a given consequence type we can estimate the number of mutations in excess for the gene-cohort in that counsequence type consequence type as:

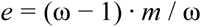

where m is the number of mutations observed with that consequence type. The aggregated excess at a gene-sample is the sum of the excess computed for missense mutations, nonsense mutations and splice-affecting mutations in that gene-sample. By adding the number of mutations in excess across the driver genes we provide an estimate of the average number of driver mutations per sample.

### BoostDM

We reported the mutation count per sample that boostDM determines to be potential drivers, which gives us a distribution of counts. For each cohort we used the most specific model that matched the gene and cohort according to the model selection strategy of boostDM (Section boostDM of Supp. Note).

We reported the distributions of counts (resp. expected counts) by boostDM (resp. dNdScv) across samples as the median and 95% confidence intervals (see Supp. Fig. S2).

### In silico saturation mutagenesis analysis

#### Classification of each possible nucleotide change

For all 105 gene tumor type specific models (section boostDM of Supp. Note) we first annotated all coding single nucleotide substitutions using cluster annotations (IntOGen), bgvep, Phylop (Pollard et al., 2010) and PhosphoSitePlus (Hornbeck et al., 2015). Briefly, bgvep retrieves the consequence type (Sequence Ontology), amino acid change and exon of the mutation in the canonical transcript using VEP.92 (McLaren et al., 2010), for any possible nucleotide substitution mapping to the canonical transcript. We then used boostDM to score the driver probability for all the mutations with consequence types accepted by the method (i.e, missense, nonsense, splice affecting and synonymous, see also Supp. Note). BoostDM models are trained using a gradient boosting approach (Chen and Guestrin, 2016). An automated pipeline to run boostDM on cancer genes across tumor types is implemented using Nextflow (Di Tommaso et al., 2017).

### Measurements and indices

For each gene we then computed two measurements: i) the “potential drivers ratio” is the ratio between the number of mutations deemed as driver (with BoostDM score greater than 0.5) and the total number of mutations in the pool of mutations for that gene (as described above); ii) the “observed-to-potential drivers ratio” estimates the ratio between the number of observed (unique) driver mutations in the sequenced samples from the tumor type and the total number of potential driver mutations.

### Mutational probability

For each gene and tumor type we computed the flat mutation rate specific for each trinucleotide context (96-channel) at each position along the CDS of the gene (CDS reference) in accordance with the mutational profile inferred by the observed mutations in non-driver genes (IntOGen) in that cohort. More specifically, from the 96-channel profile we can infer the probability p(c) that a single observed mutation belongs to context c in a sequence with balanced triplet content. We define the Mutational Probability (relative to a gene and tumor type) of a mutation with context c as follows:

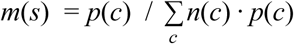

where n(c) is the number of possible mutations with context c in the gene (CDS). In other words, the Mutational Probability of a site relative to its gene is a function that maps each possible mutation site in the gene with the expected number of mutations at the site (determined by its context) conditioned to observing 1 mutation in the gene. Although the scale is relative to each gene, this expectation renders mutations comparable within genes.

### Mutational probability bias

In our analysis we were concerned with the relationship between the Mutational Probability and various ways of segregating the possible mutations in a gene. For instance, whether observed potential driver mutations tend to occur at sites with higher mutational probability compared to non-observed potential drivers.

When comparing two sets of mutations, we define the Mutational Probability Bias as the propensity that one set has higher (resp. lower) mutational probabilities than the other. For this study, this propensity is measured with two proxies: how sharply the mutational probability separates between the two groups (auROC) and what is the difference in log-scale (log-fold-change, logFC) between the median mutation probabilities of the two groups. We adopt the convention that when comparing two groups of mutations, they are labeled “True” and “False”, respectively.

We computed the area under the ROC curve (auROC) where the ROC is defined by True Positive Rate (TPR) and a False Positive Rate (FPR) for a grid of mutation probability thresholds. Specifically, in our setting the TPR are given by the ratio of “True” mutations with mutation rate above threshold over total “True” mutations; and the FPR are given by the ratio of “False” mutations with mutation rate above threshold over total “False” mutations (Fig. 4a). We also computed the log-fold-change (logFC) of the mutation rate distributions between groups as log(m/M), where m (resp. M) is the median mutational probability of the “True” (resp. “False”) group (Fig. 4a).

Interestingly, the auROC has a well-known probabilistic interpretation that the reader shall bear in mind: suppose an experiment when one sample t is randomly drawn with uniform probability from the “True” set and another sample f is drawn from the “False” set in the same way; then the probability that the t sample has higher score (e.g. mutation probability) than the f sample is precisely the auROC. The auROC is also connected to the Mann-Whitney U statistic in the sense that auROC = U / (n·m), where n, m are the sizes of the “True” and “False” groups, respectively.

In practice, we require this notion in essentially two scenarios: i) comparing observed vs non-observed mutations of some kind; ii) comparing potential drivers vs passengers.

### Signature deconstruction

The mutational spectra of the 28,076 samples in IntOGen were deconstructed into exposures of mutational signatures using the deconstructSigs package (Rosenthal et al., 2016). The set of signatures considered for the fitting was the 30 COSMIC SBS v2 (Alexandrov et al., 2013; Tate et al., 2019). Only signatures found active in the respective tissue according to COSMIC were allowed for the reconstruction (Table S3). A maximum likelihood approach (Pich et al., 2018) was used to assign the most likely etiology to each of the driver mutations. Briefly, after signature deconstruction we estimated the amount of exposure attributable to each signature for each channel (tri-nucleotide context), whence we inferred the conditional probabilities that a mutation with some context was caused by each of the mutational signatures considered to be active in the tissue. For each mutation, the signature with the maximum probability was considered as the one contributing it (maximum likelihood).

### Mutational discovery index

For each cancer gene and tumor type pair, we provided an estimation of the expected number of unique driver mutations to be found as a function of the number of sequenced samples, which we denote by E(n). This estimation gives us a way to numerically represent to what extent some driver mutations have not yet been discovered, thereby depicting the discovery status of cancer mutations per gene and tumor type. We define a continuous score, the Mutational Discovery Index, with values in the unit interval, such that higher values imply fewer expected new (non previously observed) driver mutations upon new sequencing experiments.

The mutational discovery index arises upon fitting E(n). To this end we generated a collection of data points D of the form (n, u) by picking random subsets of samples of different sizes (n, taking 20 different such values by evenly spacing the interval between 0 and the number of samples matching the tumor-type) and counting the number of unique potential drivers according to boostDM, yielding u. Additionally, for each n, 10 points were generated by picking independently with replacement subsets of size n then counting unique mutations.

The best fit *Ê*(*n*) for E(n) was computed as the best least-squares fit of the form:

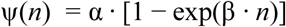

to the data points D. Justification and potential caveats of this exponential form can be found in Section “Remark on the Mutational Discovery Index” of the Supp. Note).

We formally define the mutational discovery index (MDI) of a single fit as:

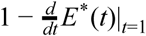

Where *E*\*(*t*) is a transform of *Ê*(*n*) to render the input and output values relative to the respective maximum values, i.e., within the unit interval [0, 1]. More specifically:

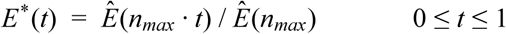

where *n*_*max*_ is the cohort size.

The MDI reported for a gene and cohort arises upon downsampling of the subsample data points (randomly selecting one point per each subsample size) and computing the median MDIs for the respective fits.

### Other software used

To carry out the analyses described in the manuscript and prepare the figures, we employed the Python programming language and several ready-to-use packages and utilities, such as Jupyter notebooks, matplotlib (Hunter, 2007), numpy (Oliphant, 2006), the pandas library (McKinney, 2017), scipy (Virtanen et al., 2020) and scikit-learn (Pedregosa et al., 2011).

## Data and Code Availability

### Data Availability

BoostDM predictions and feature explanations for all possible point mutations mapping to canonical transcripts are available for the collection of 105 genes and tumor type specific models (Fig. 1f). The results are available on the website: https://www.intogen.org/boostdm. The classification of all observed mutations in cancer genes across tumor types is also available at this site.

### Code Availability

The code is available from the following repository: https://bitbucket.org/bbglab/boostdm/

