## Supplemental Information for "In silico saturation mutagenesis of cancer genes"

1. **Supplementary Figures**

2. **Supplementary Note**

3. **Supplementary Tables**

4. **Supplementary Data**

### 1. Supplementary Figures

### Supplementary Figure 1

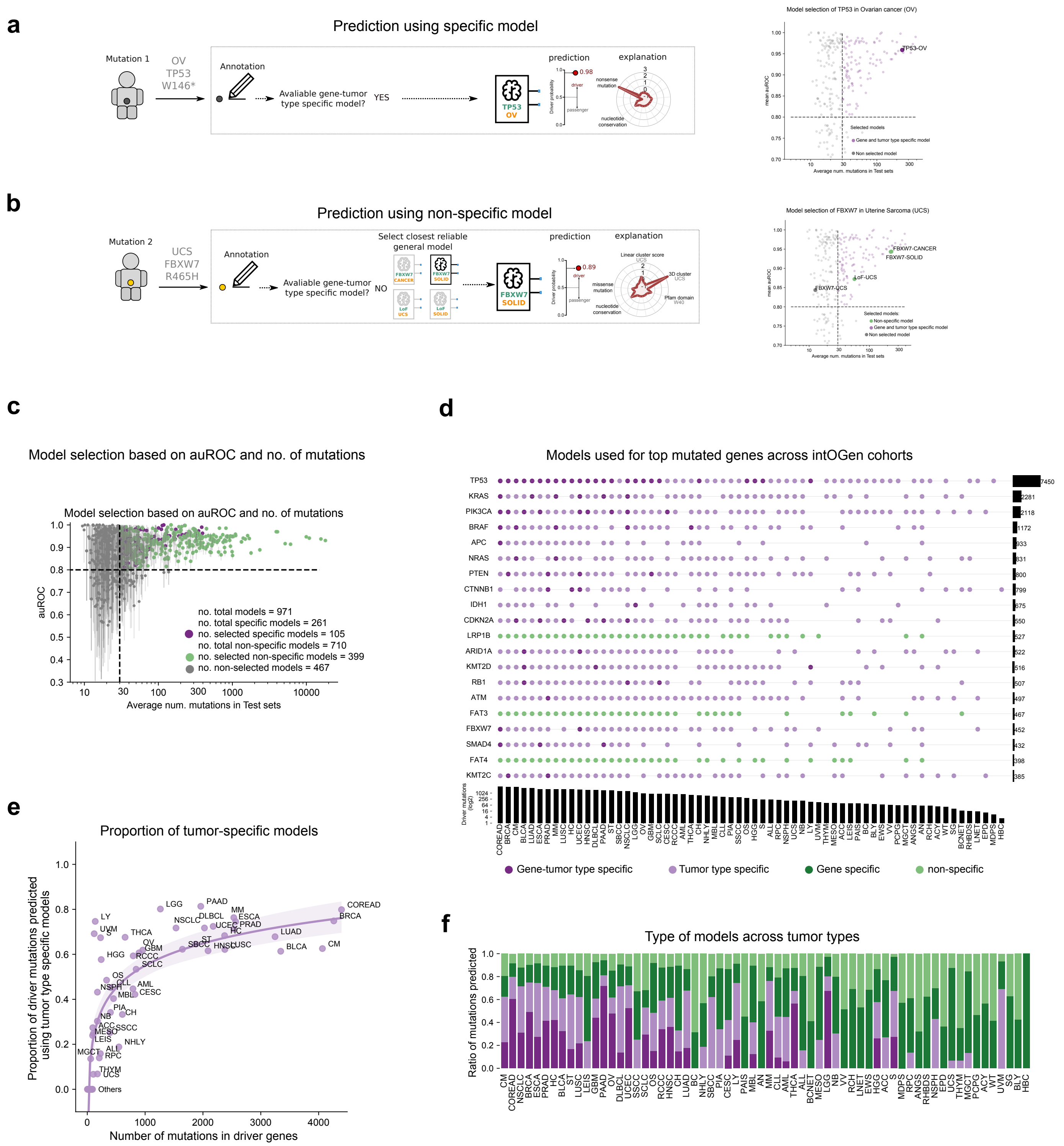

**Figure S1. BoostDM general models, model selection and results of the combination of general and specific models.**

a) In the operation of boostDM, if there is a high-quality (see main manuscript and Supp. Note) cancer gene-tumor type specific model it is used to classify the mutation in question. The application of a cancer gene-tumor type specific model is illustrated with the classification and explanation of the TP53 W146* mutation in ovarian adenocarcinomas.

b) If no high-quality model is found to classify a mutation (either due to insufficient mutations for the training or poor performance; see Supp. Note), as in the case of FBXW7 R465H mutation in uterine carcinosarcomas (UCS), boostDM searches for a more general model. Two approaches are followed to train such general models: i) pooling the mutations observed in a cancer gene across related tumor types; ii) pooling mutations observed in different cancer genes. Tumor types may be clustered following the oncotree ontology (ref; see Supp. Note) up to the root, where the ~27,000 samples of all tumor types are pooled together, while cancer genes may be agglomerated according to their mode of action. Combining these two pooling approaches, dozens of models may be built to classify any mutation. The search for the right model to classify a mutation (if a specific model is unavailable) first navigates the oncotree hierarchy, and then the mode of action of cancer genes. In the example in the Figure, a model trained on FBXW7 mutations across all solid tumors shows a performance above the threshold of reliability established for high-quality models (Supp. Note). It is thus not necessary to resort to higher-order models, such as FBXW7-cancer, Loss-of-Function (LoF)-UCS, or LoF-Solid. The scatter plot at the right of the panel summarizes the search for the best performing model to classify FBXW7 mutations in UCS.

c) Summary of the 504 models selected to classify mutations across all cancer gene-tumor type combinations in the IntOGen platform (www.intogen.org). The panel is similar to main Figure 1d.

d) Summary of the models employed to classify mutations across 20 cancer genes and 64 tumor types. The circles indicated whether a specific model, a model trained on mutations of different genes in a tumor type (tumor type specific), a model trained on mutations affecting a gene across different tumor types (gene specific), or a totally non specific model. Below the graph the total number of driver mutations identified by boostDM through the combination of all models in each tumor type are represented through a barplot. The same summary, for the driver mutations affecting each cancer gene across all tumor types, is represented at the right side of the plot.

e) Most driver mutations in cancer types with many sequenced cohorts in the public domain (www.intogen.org), such as colorectal tumors (COREAD), or breast adenocarcinomas (BRCA) are identified through specific models. These are also the tumor types with higher total number of driver mutations identified. The importance of specific models in the identification of driver mutations decreases together with the total number of drivers detected. The plot illustrates that the majority of driver mutations across cancer genes and tumor types are identified through specific models, highlighting the importance to continue analyzing cohorts of tumors that are continuously deposited in the public domain to be able to train more specific models.

f) The specific relative contribution of models of the four types defined in panel d) to the identification of driver mutations in 64 tumor types.

### Supplementary Figure 2

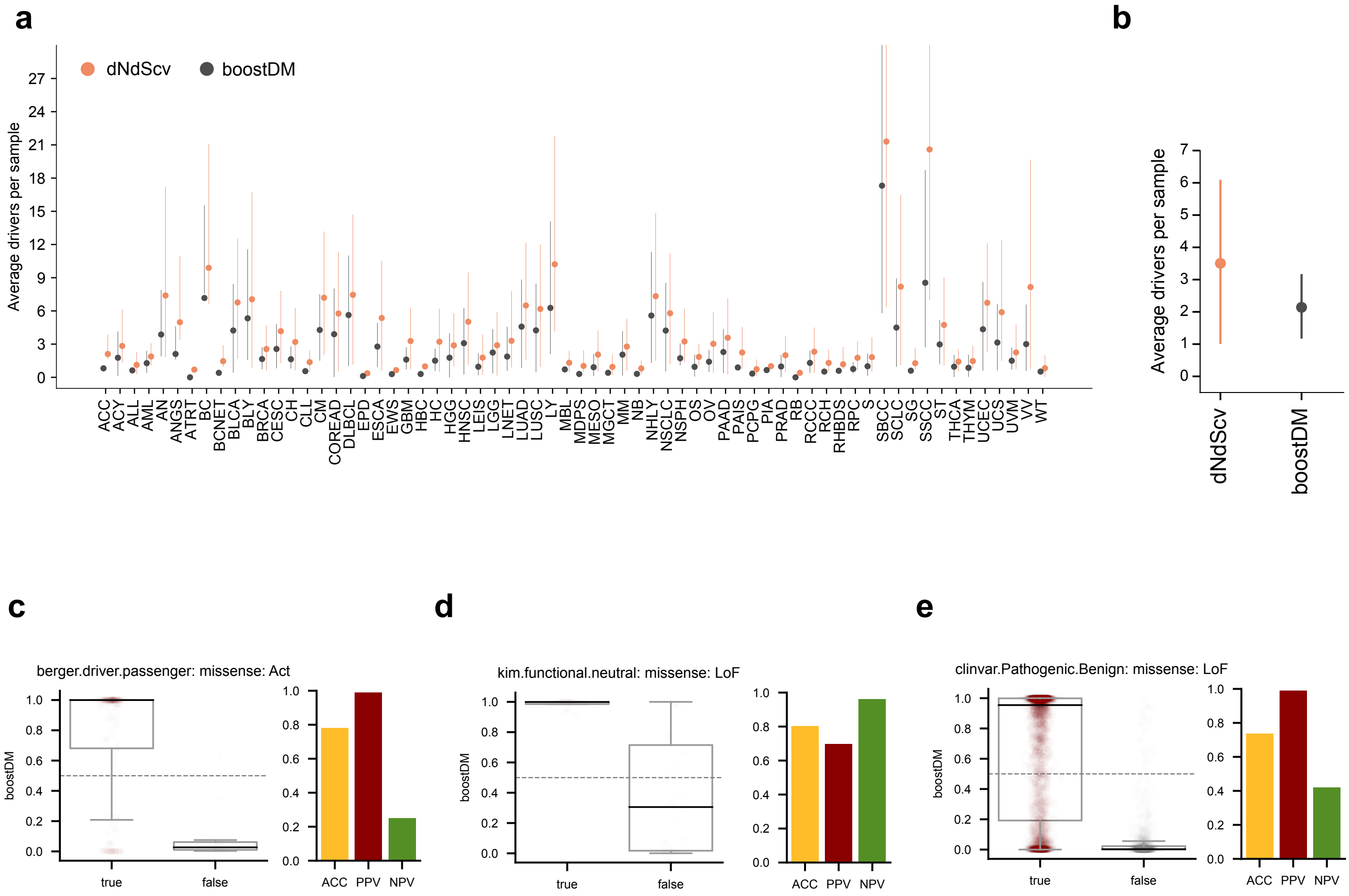

**Figure S2. Performance of boostDM in the identification of driver mutations across cancer types**

a) We compared the median (and 95% confidence intervals) of the number of driver mutations identified by boostDM (applying specific and nonspecific models) in each tumor sample of 66 cancer types with that estimated recently through the extent of positive selection (dNdScv; ref dnds). Across most cancer types, the average number of driver mutations identified through both approaches is very similar.

b) Summary of the average number of driver mutations identified by both approaches across the ~27,000 tumors of 66 cancer types (www.intogen.org).

c-e) Summary of the performance of boostDM models in the separation of experimentally validated functional and benign mutations in oncogenes in the BERGER dataset (c), functional and neutral mutations in tumor suppressor genes in the KIM dataset (d), and pathogenic and benign mutations collected from the literature in ClinVar (e). The performance is represented through the same metrics as in Figure 1h.

### Supplementary Figure 3

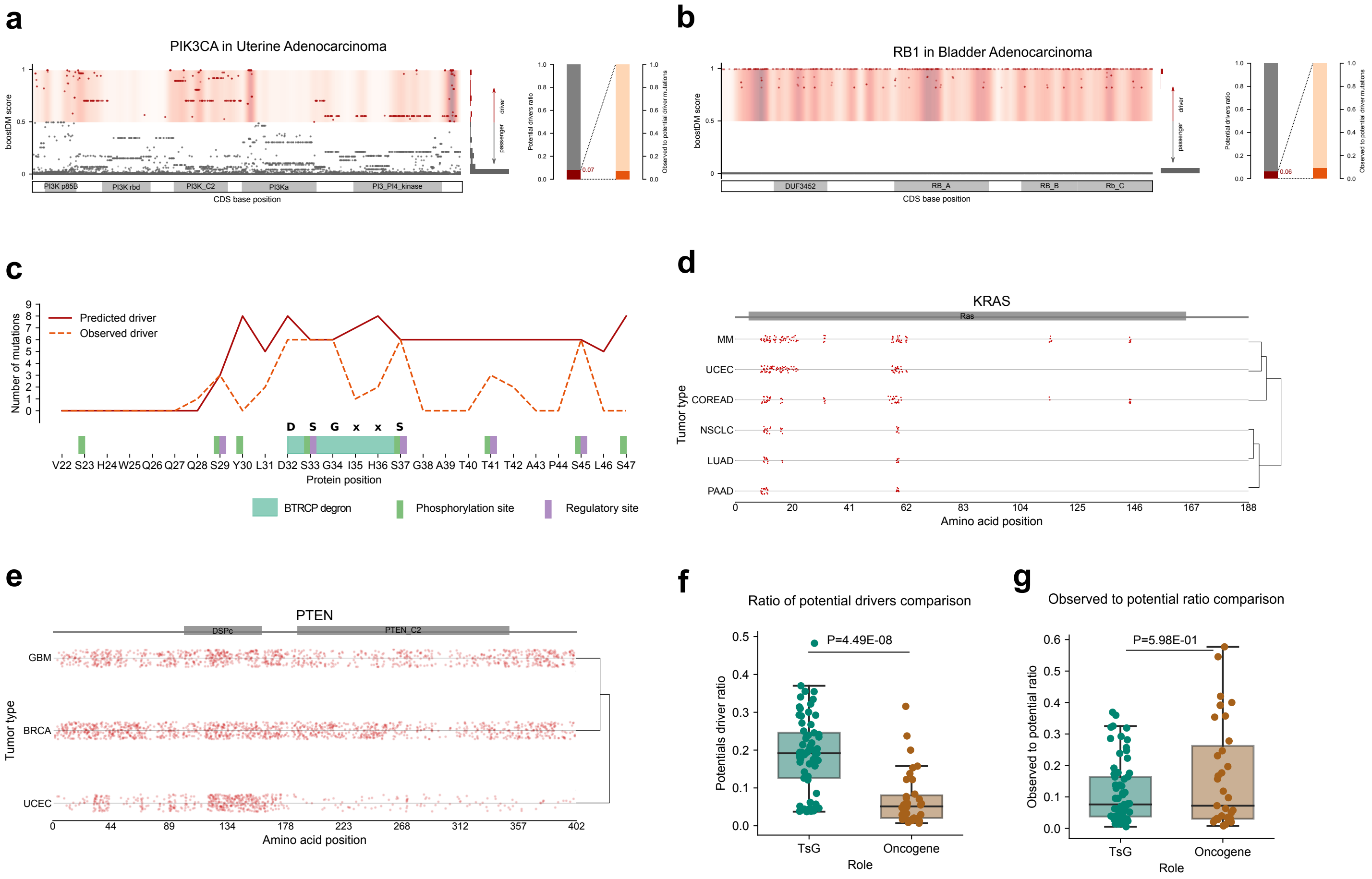

**Figure S3. In silico saturation mutagenesis of cancer genes**

a,b) Result of applying the in silico saturation mutagenesis approach to PIK3CA across uterine adenocarcinomas and RB1 across bladder tumors. Following the rationale illustrated in Figures 3a-d, all possible mutations in each cancer gene (dots) are represented in their relative position of the gene coding sequence (x-axis). Relevant functional protein domains are represented within each gene body. Potential driver mutations appear in red and potential passenger mutations, gray. The concentration of driver mutations at different regions of the protein is represented as a density. The distribution of boostDM driver scores of all possible mutations appears at the right side of each saturation mutagenesis plot. Two bar plots illustrate the potential drivers ratio (red on gray) and the ratio of observed-to-potential driver mutations (orange on gray).

c) Distribution of all (solid line) and observed (dashed line) driver mutations in and around the CTNNB1 B-TRCP degron. It zooms into this particular region of the in silico saturation mutagenesis profile of the whole gene shown in Figure 3c.

d,e) Profile of potential driver mutations of KRAS and PTEN across several tumor types.

f) The distributions of the ratio of potential drivers across tumor suppressors and oncogenes are significantly different (one-tailed Mann-Whitney test). Tumor suppressor genes show significantly greater ratios of potential driver mutations than oncogenes, since more positions are available to loss-of-function than to gain-of-function mutations.

g) On the other hand, the observed-to-potential ratio of driver mutations of both oncogenes and tumor suppressor are not significantly different (one-tailed Mann-Whitney test). Some oncogenes with particularly few positions available to gain-of-function mutations exhibit very high (above 0.5) observed-to-potential ratio of driver mutations.

### Supplementary Figure 4

**a**

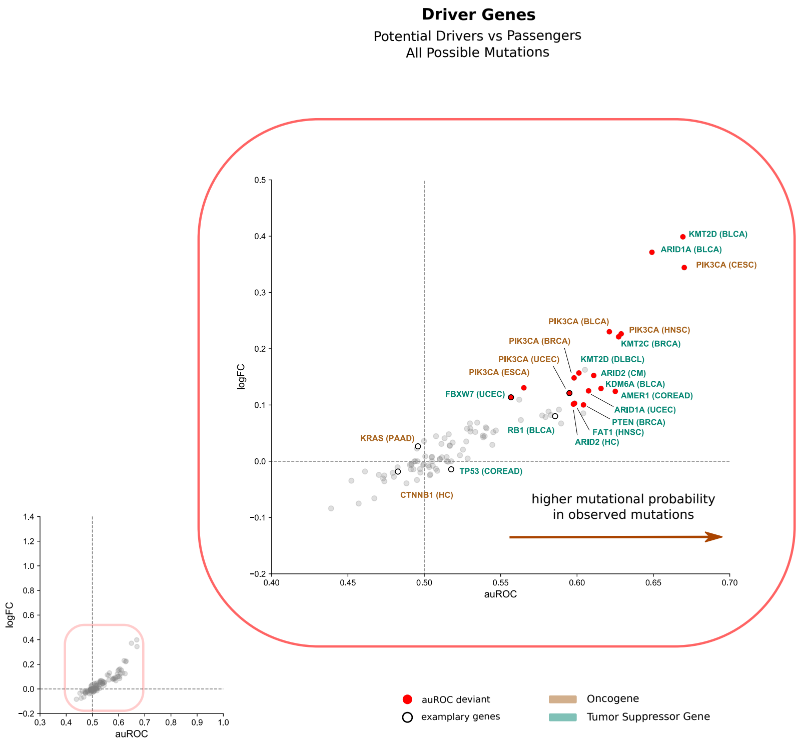

**b**

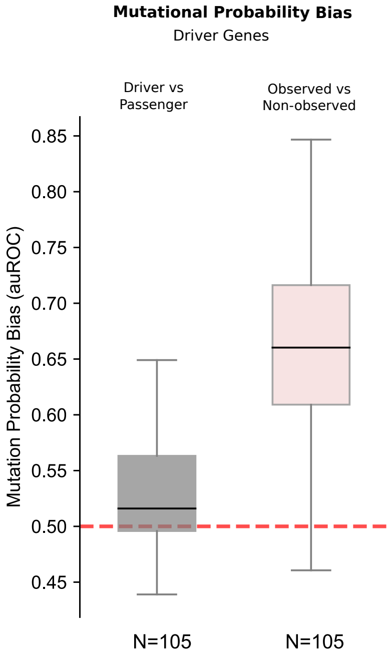

**Figure S4. Driver and passenger mutations in some cancer genes show mutation probability bias.**

We computed the mutation probability bias of driver and passenger mutations in cancer genes using the area under the ROC curve (auROC) of the separation between both sets and their log-fold change (logFC), as explained in the main manuscript and main Figure 4.

a) The distribution of mutation probability bias between driver and passenger mutations for the 105 cancer gene-tumor type combinations with specific models appears in the left bottom corner of the panel. For the majority of cancer genes, the bias is inexistent (auROC 0.5 and logFC 0). This is expected as boostDM models are designed to avoid confounding by mutation probability. On the one hand, mutational features are in general independent of the mutation probability. On the other hand, synthetic mutations used as a negative set for training are generated following the underlying probabilities from the frequencies of tri-nucleotide changes in each tumor type. Interestingly, a few cancer genes (zoom in plot inside the red square) deviate from this trend and do exhibit a mutation probability bias between driver and passenger mutations. Deviant cancer genes were identified as having logFC>0.1 and permutation test p-value<0.01 (N=1000). These deviations appear in cases in which the most explanatory mutational features are highly concordant with the mutational probability.

b) The boxplots correspond to the distribution of the mutation probability bias (auROC) between driver and passenger mutations of cancer genes (gray) presented in panel a) and the observed-to-unobserved mutation probability bias (pink) presented in Figure 4c. While the mutation probability bias between driver and passenger mutations of cancer genes is only slightly deviated from the null value (due to deviant cancer genes discussed above), the distribution of the observed-to-unobserved mutation probability bias of cancer genes exhibits a clear deviation from the null value. In the latter case (as discussed in the main manuscript), most cancer genes exhibit a clear mutational probability bias.

### Supplementary Figure 5

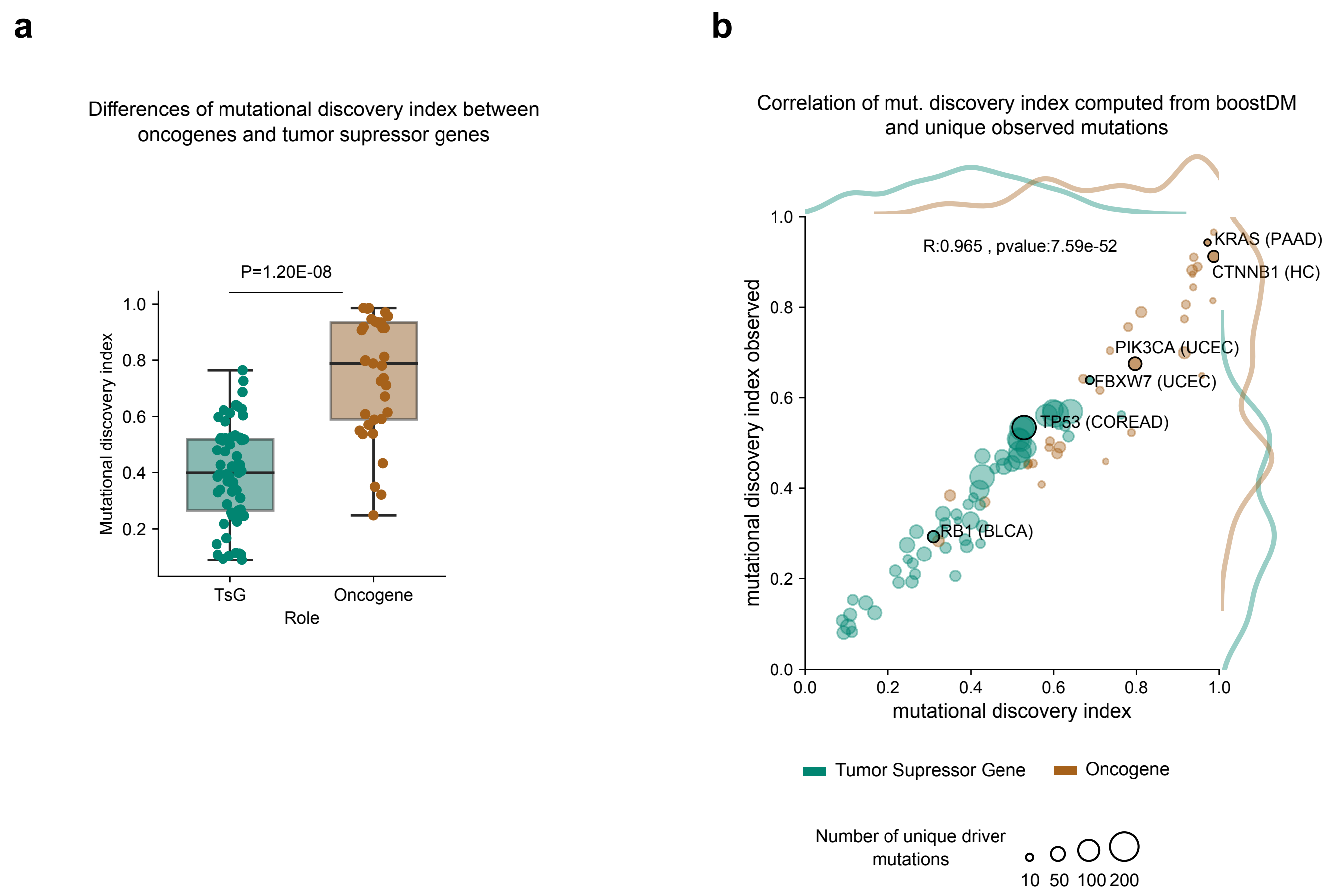

**Figure S5. Heterogeneity of mutational discovery index across cancer genes.**

a) The distributions of mutational discovery index of tumor suppressors and oncogenes are significantly different (one-tailed Mann-Whitney test). Tumor suppressor genes show significantly smaller mutational discovery index than oncogenes, since more driver mutations affecting the former are available for discovery through sequencing.

b) Two-dimensional scatter plot representing the relationship between the total number of mutations (y-axis) and the mutational discovery index (x-axis) of cancer genes with 105 specific models. Each cancer gene is represented as a circle colored by its mode of action and with size proportional to the number of unique observed mutations. The distribution of both values for tumor suppressor and oncogenes is shown along the axes. The pearson correlation coefficient representing the relationship between the two quantities and its p-value are included in the graph. This figure is similar to that shown in Figure 6g, replacing all possible mutations for all possible driver mutations. It demonstrates that the strong correlation between the number of potential driver mutations and the mutational discovery index is an intrinsic property of cancer genes, independent of the in silico saturation mutagenesis approach.

#### 2. Supplementary Note

Supplementary Note

*In silico saturation mutagenesis of cancer genes*

### 1 boostDM

We describe a method to discriminate between driver and passenger mutations in driver genes, which we have employed to conduct *in silico saturation mutagenesis*, i.e., to score all possible point mutations in cancer driver genes for their potential to be involved in tumorigenesis.

#### 1.1 Overview

`boostDM` is based on the analysis of observed mutations in sequenced tumours and their site-by-site annotation with *mutational features* encompassing relevant biological information to the selective involvement of the variant.

##### 1.1.1 Supervised learning approach

The method is based on training a supervised model using a training dataset composed of driver and passenger mutations of a cancer gene. One of the main difficulties for this is that in general there is no ground truth collection of driver and passenger point mutations in a cancer gene. However, for some genes, the observed-to-expected ratio of mutations is large enough that the vast majority of observed mutations are involved in tumorigenesis. We reason that mutations in a cancer driver gene (identified by IntOGen) above a certain excess of observed (over expected) mutations ($\geq 85\%$ estimated by dNdScv [1]) are most likely drivers and can thus be used as a positive set (Drivers) for the training. On the other hand, we convene that passengers are randomly generated mutations with flat probabilities based on the tri-nucleotide specific mutation rates recorded in the relevant tumor type. Therefore, a dataset of synthetic mutations generated following these probabilities can be used as a negative set (Passengers).

##### 1.1.2 Features

Each mutation provided for training, in both Drivers and Passengers sets, is annotated with relevant mutational features, which the classification task exploits to discriminate between observed drivers and passengers in tumours.

Mutational features of each cancer gene across malignancies have been derived from the systematic analysis of tens of thousands of tumor samples (IntOGen [2]). Other relevant features are collected from public databases of biological sequences (see Key Resources Table in Methods).

##### 1.1.3 Ensemble of classifiers

For the sake of preventing overfitting, several classifiers are trained in parallel with random partial subsets of the training data, giving rise to a pool of expert classifiers that can be merged into a consensus forecast. Alongside the training, the method also yields a cross-validation assessment of the performance of the models.

#### 1.2 Scope

`boostDM` looks into the protein coding sequence of the genome. All mutations in the training sets map to the canonical transcript of protein coding genes according to the Ensembl Variant Effect Predictor version 92 (VEP.92 [3]). Notice that these transcripts may include mutations in untranslated regions (UTRs) and splicing affecting mutations, thus non-protein-altering sites.

#### 1.3 Prediction and Explanation

For each cancer gene and tumor type our method constructs a model that represents the combinations of features that define driver mutations in that gene and cancer type. Given the context and the features for a mutation, the method yields a score $p \in [0, 1]$ in the unit interval (`boostDM` score) that reflects the strength of the forecast that the mutation is involved in tumorigenesis (potential driver): higher `boostDM` score implies stronger evidence of driver potential. Although the final score is not calibrated to support a purely probabilistic interpretation, by design a score $> 0.5$ is to be interpreted as reflecting positive evidence to deem the mutation a potential driver.

Moreover, the supervised learning approach further allows breakdown of the forecast of individual mutations in terms of the so-called SHAP values [4]. Intuitively, these are explanatory values in the sense that if a feature has positive (resp. negative) value, then the method deems more likely (resp. unlikely) that the mutation is a driver given the value of the feature in relation to the other features, i.e., the feature contribution for that individual prediction is more decisive (see also Section 1.9).

#### 1.4 Models

##### 1.4.1 Specification

Each `boostDM` model is an ensemble of expert classifiers ($n = 50$) each trained with a partial view of the training dataset. Each expert classifier is a boosted trees model (sum of tree functions fitted by gradient boosting) with a logistic binary objective function (cross-entropy loss). The individual predictions are merged into the `boostDM` score with an aggregator function of classifiers intended to correct for the systematic bias of each expert classifier (Section 1.10). The reported explanations are the average of the SHAP values yielded by all expert classifiers (Section 1.9).

##### 1.4.2 Implementation

The `boostDM` pipeline has been implemented in Python [5] and Nextflow [6]. The key ingredients for our models are the libraries `xgboost` [7] (version 0.90) for training the gradient boosting classifiers and `shap` [4] (version 0.28.5) for the computation of the SHAP values associated with the predictions. A complete list of dependencies is given in the main Methods (Resources).

##### 1.4.3 Hyperparameters

Intuitively, the model hyperparameters specify the learning strategy by which the optimal tree function is searched for in order to keep a good balance between minimization of the objective loss and prospects for good generalization. While the values of some hyperparameters are the result of an explicit design decision, others are not unequivocally defined. The current values stand as an engineering compromise resulting from repeated testing and refining. For more information about the gradient boosting models used, check `xgboost` documentation: xgboost.readthedocs.io.

Herein we provide the main hyperparameter settings alongside their meaning: i) The model comprises sums of binary tree functions (`booster = "gbtree"`). ii) The learning task minimizes a cross-entropy loss function (`objective = "binary:logistic"`). iii) All the features are available when building each new tree (`colsample_bytree = 1`) and each new tree level (`colsample_bylevel = 1`). This was deemed necessary as `boostDM` uses a small set of features. iv) Learning rate 0.001 (`learning_rate = 0.001`) was decided on by testing. v) Percentage of samples randomly drawn prior to growing a new tree is 70% at every iteration (`subsample = 0.7`). This is a technical choice to prevent overfitting decided on by testing. vi) Maximum depth of trees used in the tree function is 4 (`max_depth=4`). In practice a good performance can be attained even at depth as low as 1 (stumps). However, at a higher computational cost, more depth gives also more room for learning complex dependencies. Our choice was decided on by testing. vii) Default regularization hyperparameters were employed. viii) Maximum number of training steps is 20,000 (`n_estimators = 20000`).

#### 1.5 Ontologies

We aim to develop models that are capable of classifying all mutations across all driver genes and tumor types. The tumor type value determines the set of sequenced samples that provide the set of observed mutations and the trinucleotide based probabilities to generate synthetic mutations. In order to properly index the models (i.e., annotate to which cancer genes and tumor types they are applicable) we resort to two ontologies for gene and tumor-types. Importantly, these ontologies will comprise terms with several degrees of specificity, so that we can implement a flexible model selection strategy to extend the classification to all cancer genes in the compendium of mutational drivers across tumor types (see Section 1.11).

For genes we will consider a simple hierarchy whereby a root term "GENE" has two children "LoF" (tumor suppressor) and "Act" (oncogene), which in turn have gene names (gene identifier) as children, according to their mode of action (derived from IntOGen). Those genes that are labeled with "Ambiguous" mode of action have "GENE" as parent. We will refer to this hierarchy as Gene Hierarchy.

We employ a simplified tumor type ontology, referred to as the Oncotree, adapted from IntOGen [2]. This ontology allows us to group samples according to several degrees of specificity regarding the tissue-pathology context where the mutations are reported. Thus, a root term “CANCER” is connected to two children terms, “SOLID” and “NON-SOLID”, from which new children terms arise at increasing specificity. The leaves of this hierarchy define the most specific tumor-type terms considered in this study (see Table S2 and Figure SN1 below):

#### 1.6 Index of Models

Each model is labelled with a unique pair of gene and tumor-type terms, i.e., each model is to be understood in relation to a particular cancer gene and a tumor-type term in the Oncotree. We record this convention in the notation by writing (G,T)-classifier or (G,T)-model when referring to an expert classifier (resp. model) indexed by gene G and tumor-type T.

#### 1.7 Data Processing

##### 1.7.1 Source

The following essential data inputs were collected from IntOGen [2]: the compendium of 568 mutational driver genes alongside their driver discovery output annotations, including consequence-type-specific dN/dS (dNdScv) and mode of action per gene; the catalogue of observed mutations in those genes; site-specific mutational features associated with various signals of positive selection, including: 3D clusters, linear clusters and recurrently mutated domains.

##### 1.7.2 Filtering

**Consequence type**. We restricted our analysis to point mutations with either of the following Sequence Ontology [8] terms (according to VEP.92): splice-donor-variant, splice-acceptor-variant, splice-region-variant; missense-variant; stop-gained; synonymous-variant.

**Multiple nucleotide variants**. Adjacent point mutations were excluded from our analysis due to the plausible risk that these are misannotated multiple nucleotide variants.

##### 1.7.3 Mutational features

Given a point mutation in a driver gene and tumor type (either observed or randomized) our method requires an annotation of the mutation with a series of features to train the classifiers.

- Consequence type: every mutation was annotated with the following binary features, matching the obvious Sequence Ontology annotations: `missense`, `nonsense`, `synonymous` and `splicing`. For all observed and synthetic mutations in driver genes, we retrieved the annotations from VEP.92 for the canonical transcript (see Section 1.7.2).
- Clusters of mutations in the DNA primary structure (linear clusters): for every mutation we annotated whether it overlaps a significant linear cluster identified by the method OncodriveCLUSTL [9]. We created two annotation tiers for mutations overlapping i) linear clusters found in a cohort of the corresponding tumor type (`cat_1`), or ii) clusters only identified in other tumor types (`cat_2`). Additionally we create another feature that represents the OncodriveCLUSTL score of the linear cluster in the tumor type (i.e., `cat_1`).
- Clusters of mutations on the protein 3D structure (3D clusters) identified by the method HotMAPS [10] in a tumor type specific (`cat_1`) or pan-cancer (`cat_2`) manner. For details of the HotMAPS implementation please refer to [2].
- The overlap with Pfam domains [11] that are significantly enriched for mutations in the gene across tumors of the cancer type, identified by the method smRegions [12].
- Phylogenetic conservation of the nucleotide across mammals measured through the PhyloP score [13].
- Non-synonymous coding mutations were also annotated with post translational modifications (PTMs) when the mutation affected an amino acid

that is known to be acetylated, phosphorylated, ubiquitinated, methylated or subjected to any other regulatory modification according to PhosphositePlus [14].

- Nonsense mutations with a special annotation stating whether or not they overlap the last coding exon of the canonical transcript (according to VEP.92) reflecting the potential for the truncating variant to skip non-sense RNA-mediated decay.

- Mode of action of the driver gene harbouring the mutation, either activating (Act), loss-of-function (LoF) or Ambiguous. This feature is only relevant for general models involving mutations across all driver genes (see Model Selection).

Nonsynonymous point mutations were annotated with linear clusters, 3D clusters or Pfam domains if the genomic coordinate of the mutation overlapped a significant linear cluster (resp. 3D cluster or Pfam domain) according to IntOGen across any cohort of the corresponding tumor type (or any tumor type in the case of the pan-cancer annotation). Nonsense mutations included an annotation stating whether they overlap the last coding exon of the canonical transcript according to VEP.92. Finally, all SNVs included an annotation of the affected nucleotide conservation, i.e., their PhyloP score.

Two features (PhyloP and the OncodriveCLUSTL score) were encoded as their original floating number. The rest of the features (categorical) were encoded with a one-hot encoding.

#### 1.8 Training

##### 1.8.1 Training datasets

The first requirement for our supervised learning approach is to create a catalogue of mutations labeled as Drivers or Passengers. This catalogue is established globally for all the models, then for the training of each (G,T)-classifier only the mutations relevant to the (G,T) context are used (see Section 1.6 for notation).

##### 1.8.2 Drivers

The set of Drivers used for training are observed mutations in mutational cancer genes (IntOGen) that exhibit a consequence-type specific excess of observed (over expected) mutations above 85% (according to the method dNdScv [1]). Repeated mutations (i.e., when the same mutation is observed in different samples) are allowed in the set of Drivers. The choice of the excess is a trade-off between the confidence that the mutations included in Drivers play a real driver role and the amount of genes for which this training can be carried out.

##### 1.8.3 Passengers

For each set of Driver labels a comparable set of Passenger labels is generated. For each Driver mutation observed in a gene and cohort of samples, we generate 50 randomly and independently chosen mutations with replacement in the gene (VEP.92 canonical transcript), with probability proportional to the average site-specific mutation rate recorded in the most specific tumor-type (Oncotree) matching the cohort (see Methods).

##### 1.8.4 Data Splits

A (G,T)-classifier results from training a gradient boosting classifier with two sets of annotated mutations: Train and Test. We will refer to a Train-Test pair as a Split. In our setting Splits are randomly generated and must satisfy the following conditions:

1. Both, Train and Test are balanced sets, i.e., in each set there is the same number of Driver and Passenger labels.
2. The sizes of Train and Test are in a 70:30 ratio (70/30 cross validation).
3. Each Driver mutation observed in the (G,T) context appears only once in either Train or Test.
4. Repeated Driver mutations (i.e., when the same mutation is observed in different samples) are allowed in Train, but not in Test, in order to prevent spurious inflation of the cross-validation performance evaluation.

5. Passenger mutations in Train and Test are randomly selected among all the mutations in the Passengers pool matching the (G,T) context.

Each Split completely determines a model fit given the set of hyperparameters, i.e., the learning configuration of the learning algorithm (see Section 1.4.3)

##### 1.8.5 Cross-validation and early stopping

When training each classifier, we implement a cross-validation tactic to prevent overfitting, consisting of evaluating the performance of a partial model trained with the Train dataset after each learning step on the Test dataset (generically referred to as cross-validation). This sequential evaluation informs the training progress after each step, in particular whether the training must stop due to steady or reduced performance for a set of consecutive iterations (early stopping).

We use the log-loss (see `scikit-learn.org/model_evaluation`) as a performance measure to assess training progress with cross validation. Given true labels $\mathbf{y} = y_i$ and forecasts $\hat{\mathbf{y}} = \hat{y}_i$, the log-loss is defined as:

$$J(\mathbf{y}, \hat{\mathbf{y}}) = -\frac{1}{N}\sum_{i=1}^{N}\left[y_i \log \hat{y}_i + (1 - y_i)\log(1 - \hat{y}_i)\right].$$

We define an early-stopping requirement of 2,000 iterations. Thus each expert classifier training finishes whenever either of the following conditions is first met: i) the maximum number of training steps (N=20,000 iterations) is attained; ii) the model's performance on the Test data set does not improve for 2,000 consecutive iterations. See Figure SN2.

After the training we record the performance of the classifier via log-loss and area under the ROC (Receiver Operating Characteristic) when applied to the Test dataset. Although cross-validation and early stopping allow some extent of data leakage from the Test dataset, it gives us a first line of model performance evaluation which is used for model selection.

#### 1.9 Explanations

Each expert classifier also yields an additive explanation model based on Shapley Additive Explanations (SHAP values) [4]. Specifically, each expert classifier can decompose the logit forecast yielded by a particular mutation $z$, say $\mathrm{logit}(p_z)$ into a collection of SHAP values $\{s_i(z)\}$, one per feature, in such a way that:

$$s_1(z) + \ldots + s_n(z) = \mathrm{logit}(p_z) \text{ for all } z.$$

The intuition behind the concept of SHAP value is rooted and better understood in the context of cooperative game theory, where the Shapley value [15] was originally discovered. If we think of $\mathrm{logit}(p)$ as a utility function that depends on the contribution of a coalition of features (thought of as players of a cooperative game) the SHAP values account for the relative contributions of the features by averaging the differences between two expectations: i) the expected forecast when the feature value is known; ii) the expected forecast when the feature value is not known.

More specifically, given an individual expert classifier, denoted by $M$, the additive explanation model $\mathcal{A}(M)$ for $M$ is a map from the feature space $F = F_1 \times \ldots \times F_n$ (the set of arrays representing the possible values of the features) onto the euclidean space $E = \mathbb{R}^n$ of dimension equal to the number of encoded features, which verifies the following two requirements:

1. The additive explanation model $\mathcal{A}(M)$ gives an additive decomposition of the (logit) forecast given by $M$, i.e. if $z \in F$ denotes the array of feature values for some individual mutation and $M(z)$ denotes the (logit) prediction of the classifier on $z$, if

   $$\mathcal{A}(M)(z) = (\phi_1, \ldots, \phi_n)$$

   then

   $$M(z) = \phi_1 + \ldots + \phi_n.$$

2. Given a feature array $z$ for some individual mutation, the $i$-th component of $\mathcal{A}(M)$ is an estimate of the average marginal payoff of the $i$-th feature over all possible feature coalitions, i.e.,

   $$\phi_i(z) = \sum_{S \subseteq [n] \setminus \{i\}} K(S) \cdot (f(S \cup \{i\}; z) - f(S; z))$$

where the sum runs through all possible subsets of the set of features excluding the $i$-th feature; and

$$K(S) = \frac{|S|! \cdot (n - |S|! - 1)}{n!}$$

is an averaging coefficient that depends on the size of S; and

$$f(U; z) = \mathrm{E}(M(z) \mid z \in U)$$

is the conditional expectation of the model's prediction for $z$, given that only the values of the features belonging to the subset $U$ are known.

#### 1.10 Consensus of expert classifiers

One potential caveat of our approach is that during training some Driver labels may correspond to mutations not involved in tumorigenesis. This implies that each individual expert classifier may be at risk of being biased by the specific choices made in the Splits. To work around these difficulties, we propose to use a pooling of classifiers, each trained with a different partial view of the data, so that given a mutation the forecast is achieved by combining the forecasts of the individual classifiers.

In the interest of clarity and interpretability of the outcome, we require `boostDM` forecasts to be sharp, i.e., with pooled probability values close to either 0 or 1. Our approach resorts to a non-linear combination of probabilities based on a logit-normal model [16]. Specifically, if a classifier $M_i$ casts a prediction $p_i$ that a specific mutation is a driver and $Y_i = \mathrm{logit}(p_i)$, this modelling choice assumes that $p_i$ arises from some latent true probability $p$ that the mutation is a driver, which has in turn been moderated by some degree of systematic bias:

$$Y_i = \log \left( \frac{p}{1-p} \right)^{1/a} + \epsilon_i$$

with $\epsilon_i \sim \mathrm{N}(0, \sigma^2)$ being a normal random variable with standard deviation $\sigma$ and $a \geq 0$ represents the amount of systematic bias.

Intuitively, systematic bias quantifies the extent to which individual classifiers regress towards a log-odds zero due to partial information or under-confidence while issuing their respective forecasts. While $a = 1$ would be associated with an accurate forecast, $a > 1$ represents under-confidence.

Using the previous modelling assumptions, given a collection of classifiers and their respective forecasts $p_i$ and logits $Y_i$, the latent true probability $p$ can be estimated as $\hat{p}$ with the following estimator:

$$\hat{p}(a) = \frac{\exp a\overline{Y}}{1 + \exp a\overline{Y}}$$

where $\overline{Y}$ denotes the average of the logits $\{Y_i\}$ and $a$ is the systematic bias.

In practice, the level of systematic bias gives us a method to sharpen the consensus forecasts of under-confident predictors. Upon testing the method in some exemplary (G,T) contexts with an MLE estimation of systematic bias [16], we committed to the choice $a = 2.3$ as uniform systematic bias.

##### 1.10.1 Model Quality Criteria

As a principle, for each (G,T) context we can compute a classification model as an aggregator of expert classifiers. However, in practice, for some models with few mutations in the training set the performance (computed as the auROCs on the respective Test datasets) may be poor.

A weak predictive power can come about because there are not enough observed mutations to render a clear signal by means of regression with the current features or because regardless of the number of mutations the set of mutational features is not sufficient to learn a distinct signal for the Drivers (see section 1.7.3).

Conversely, even if an expert classifier renders a high auROC, if there are few unique mutations in the training set the classifier may render spurious separation just by chance. This is reflected in the comparably higher spread of the auROCs in classifiers trained by less mutations (see main Fig. 1 and Supp. Fig. 1).

In view of these remarks, we establish the following quality criteria for a model to have an acceptable performance: i) mean auROCs of the constituent classifiers $\geq 0.8$; ii) mean size of the Test sets of the training Splits in the (G,T) context $\geq 30$. Notice that this implies to have at least 50 observed mutations in the (G,T) context, although this is not a sufficient condition.

#### 1.11 Selection strategy

Consider mutations in a driver gene G across tumors of cohort $C$ (according to IntOGen). What model should be used to cast a prediction on these mutations? Herein we propose a model selection rule to address this question.

For this approach we will have to bear in mind the two ontologies needed to specify indices for the models: the Gene Hierarchy and the Oncotree (see section 1.5).

For the sake of clarity in defining the selection strategy, let us introduce a minimum of notation. We denote $\mathcal{T}(C)$ the most specific tumor type term in the Oncotree matching cohort $C$, i.e., $\mathcal{T}(C)$ matches $C$ and it has no children in the Oncotree. Given any term $X$ in the Oncotree (resp. Gene Hierarchy), we denote the (unique) parent term of $X$ as $\mathrm{Par}(X)$.

##### 1.11.1 Base case: Specific Models

If the (G, $\mathcal{T}(C)$)-model meets the quality criteria defined above, then this model will be used to classify mutations in gene G across cohort $C$.

##### 1.11.2 General case: Climbing the Model Index

Now assume that the (G, $\mathcal{T}(C)$)-model does not meet the quality criteria, and thus, we decide not to use it. The following rule is intended to scan the gene-tumor-type hierarchies in order provide the best available model that reflects maximum specificity at gene level, then at tumor-type level:

1. If a candidate (G,T)-model does not meet the quality criteria, the next candidate to be checked should be the (G, Par(T))-model, given that Par(T) exists. The only exception being when T="CANCER".
2. If (G, "CANCER") is a candidate and does not meet the quality criteria, the next candidate is the (parent(G), $\mathcal{T}(C)$)-model, given that Par(G) exists. The only exception being when G="GENE".
3. If no candidate model is generated in this way meets the quality criteria, then we convene that the ("GENE", "CANCER")-model applies.

An example of how these rules apply is showed in Figure SN3.

### 2 Benchmark

We conducted validation analyses with independent experimental saturation mutagenesis assays and manually curated datasets to assess the ability of `boostDM` to identify potential driver mutations. While multiple studies have relied on in-vitro and in-vivo studies to define datasets of mutations with this capacity, the discrepancies in the definition of driverness alongside the heterogeneous experimental settings prompted us to evaluate our methodology with an array of different datasets.

#### 2.1 Experimental Saturation Mutagenesis

##### 2.1.1 Datasets

TP53 data was obtained from [17]. In the experiment, a site-directed mutagenesis was employed to construct all possible missense substitutions in the p53 protein (2569 nucleotide substitutions, 2314 different amino acid variants), followed by a promoter-specific transcriptional activity in yeast-based functional assays. Non functional sites were considered when less than the 50% wildtype transactivation was observed, as measured by the median of WAF1, MDM2, BAX, h1433s, AIP1, GADD45, NOXA and P53R2 factors.

PTEN data was obtained from [18]. The authors used a parallel approach in an artificial humanized yeast model to evaluate the impact of 7244 amino acid PTEN variants in the lipid phosphatase activity. The final score is defined by the mean of the functional score defined by the authors out of six experiments (two biological replicates with three technical replicates each).

##### 2.1.2 Analysis

We assessed the concordance between the functional scores and the binary classification produced by `boostDM` across missense mutations in all tumor-type contexts where a gene-specific model is available: for each model we computed the auROC score, where the ROC is given by: True Positive Rate (TPR) = `boostDM` positive with functional score above t over `boostDM` positive; and False Positive Rate (FPR) = `boostDM` negative with functional score above t over `boostDM` negative; with running threshold t. The experimental mutagenesis functional score separates drivers and passengers with all TP53 models with auROC above 0.75 (Fig. 1e). The same comparisons in PTEN renders a less sharp separation (see Figure SN4).

##### 2.1.3 Discussion

Saturation mutagenesis experiments are important tools to understand the functionality of all possible mutations affecting a cancer gene and their potential relevance in tumorigenesis. Nevertheless, several caveats of the experimental setting must be taken into account in this reasoning. First, most of these experiments carry out some measure of functional impact of possible mutations in the gene under study. How much the functional impact of a mutation reflects its potential tumorigenic capability is something to be determined and depends, among other things, on the mode of action of the cancer gene. For example, high impacting mutations in tumor suppressor genes are likely to be tumorigenic, but identifying gain-of-function mutations in oncogenes is arguably more difficult. Another problem is that saturation mutagenesis experiments reflect the biology of the cell type (or the in vitro construct) where they have been assayed. How the functional impact of mutations studied in a particular setting translates into a tumorigenic capability across different cell types with varying biology is also difficult to assess.

Our in silico saturation mutagenesis based on `boostDM`, on the other hand directly evaluates the likelihood of each possible mutation of a cancer gene to be involved in tumorigenesis. This likelihood is based on the features that delineate observed driver mutations and are thus likely to provide a more direct assessment of the tumorigenic potential of a mutation than the functional impact computed through saturation mutagenesis experiments. Furthermore, we construct tumor type-specific models that are able to capture the particularities of driver mutations of a cancer gene in different tissues. In summary, these differences between our silico saturation mutagenesis approach and experimental saturation mutagenesis experiments may explain the divergence of the results of both for certain cancer genes.

#### 2.2 Validated Cancer Related Variants

##### 2.2.1 Datasets

Rare Oncogenic Variants (KIM). In [19] the authors validated a set of cancer mutations which comprises both high and low frequently observed mutations through in vivo tumor formation assays. Out of the positive set, 71 alleles belonging to the rarely observed category were validated by xenografts in NCR-Nu mice, as measured by the size of the tumors after 130 days. We used the "functional" (set of positive) and "neutral" (set of negative) labels as defined by the authors.

ClinVar [20] is a database that annotates human variation with phenotypic consequence. Clinvar annotated vcf was downloaded on 2nd of March 2020. Mutations labeled as "Pathogenic" or "Likely Pathogenic" in the somatic category were used as "Driver". Mutations labeled as "Benign" or "Likely Benign" in the germline category were used as "Passenger".

High-throughput phenotyping of somatic mutations in lung cancer (BERGER). In [21] the authors experimentally analyzed 194 mutations observed in lung adenocarcinomas, 69% of which were deemed as impactful via expression-based variant impact phenotyping (eVIP). Resistance to EGFR inhibition by erlotinib in two different concentrations was also measured. We labeled the mutation as "Driver" when an eVIP-positive mutation also elicited resistance to EGFR inhibition in both conditions, and "Passenger" when the allele was negative in both experiments.

OncoKB [22] is a database that annotates the oncogenicity of several somatic mutations. We selected the "oncogenic" labeled mutations as a "Driver" set.

##### 2.2.2 Analysis

The annotations in KIM, ClinVar and BERGER datasets render a collection of mutations with binary "Driver" or "Passenger" labels. We assessed the concordance between the binary classification given by the respective datasets and the `boostDM` classification. We segregated mutations by mode of action of the respective genes and consequence-type of the mutation. For each pool of mutations we computed the respective confusion matrix, whence the positive concordance scores (resp. negative) predictive value (PPV and NPV) and accuracy (ACC) were computed. Additionally, concordance scores were also computed gene-by-gene for genes where at least one "Driver" and one "Passenger" annotated mutations were found.

### 3 Remark on the Mutational Discovery Index

For the sake of completeness, we provide justification for the exponential function used to define the Mutational Discovery Index.

#### 3.1 An Elementary Urn Problem

Consider an urn containing N different mutations that can occur in some gene. Each mutation has in addition one of two colors: driver or passenger. There are $d$ driver and $p$ passenger mutations, hence $N = p + d$. If we sample $n$ times with uniform probability and with replacement one mutation at a time from the urn, *what is the expected number of driver mutations drawn at least one time?*

This problem is related to the more general problem of estimating the probability of observing a not-yet-observed instance, given a series of past observations of instances.

Let the solution of our problem be $E(n)$, a function that maps the number of times we draw from the urn to the expected number of unique driver mutations drawn. There are two trivial base cases: $E(0) = 0$ and $E(1) = d/N$. For the general case $n > 1$ observe that we can use the following recurrence:

$$E(n) = E(n-1) + \frac{1}{n} \cdot (d - E(n-1)) = \frac{N-1}{N} \cdot E(n-1) + \frac{d}{N}$$

namely, the expectation at step $n$ must be the expectation at step $n-1$, plus the probability to draw a not previously drawn mutation in the last draw. If we let $a = (N-1)/N$, this recurrence leads us to the following expression:

$$E(n) = \frac{d}{N} \cdot \frac{1-a^n}{1-a} = d \cdot \left[1 - \left(\frac{N-1}{N}\right)^n\right]$$

Now suppose that the probability $\delta$ to draw any particular driver is uniform across drivers, but not necessarily equal to $1/N$. We can build a similar recurrence:

$$E(n) = E(n-1) + \delta \cdot (d - E(n-1)) = (1-\delta) \cdot E(n-1) + \delta \cdot d$$

Whence the following expression if we let $a = 1 - \delta$:

$$E(n) = \delta \cdot d \cdot \frac{1-a^n}{1-a} = d \cdot [1 - (1-\delta)^n] .$$

For $d$ small, notice that we have the approximation:

$$E(n) \approx d \cdot [1 - \exp(-\delta \cdot n)]$$

All in all, the previous expressions can be both put into exponential form:

$$E(n) = d \cdot [1 - \exp(\beta \cdot n)]$$

where $\beta$ is either $\log((N-1)/N)$ or $\log(1-d)$, depending on the case. This is the expression we used to define an index measuring the extent to which more driver mutations are to be discovered provided more sequenced tumors. Intuitively, $d$ is the asymptotic value that the function $E(n)$ would take for $n$ large, i.e., the total number of driver mutations to be discovered in the Urn. Notice also that the smaller is $\beta$ the faster the exponential term approaches zero, i.e., the fewer samples are required to approach the asymptotic value.

#### 3.2 Discussion

The previous assumptions are too simplistic to match exactly a real scenario, as probabilities of driver mutations to show up in a sequencing experiment may in general not be uniform. In fact in some tumor-types some driver mutations are very common, while others are only rarely observed.

Our choice can be justified for the practical purposes in our analysis. First, in many cases our exponentials seem to adjust well the data, with no remarkable pathologies showing up. Second, by straightforward enrichment of the assumptions to imply the more general case with many driver probabilities, e.g. multiexponential fitting, the method would easily bear methodological caveats itself, e.g. the requirement of good prior estimates.

Estimation of the probabilities to observe instances of not-yet-observed mutations given the observed ones, alongside accurate estimation of their prevalence across cohorts of sequenced tumours, is really the key to yield a more robust solution. Different strategies have been proposed based on the well-known Good-Turing statistic, some adapted to the problem of forecasting rare somatic variants e.g. [23]. We acknowledge that a deeper understanding of the unobserved driver landscape may bear these techniques.

[22] D. Chakravarty, J. Gao, S. M. Phillips, R. Kundra, H. Zhang, J. Wang, J. E. Rudolph, R. Yaeger, T. Soumerai, M. H. Nissan, M. T. Chang, S. Chandarlapaty, T. A. Traina, P. K. Paik, A. L. Ho, F. M. Hantash, A. Grupe, S. S. Baxi, M. K. Callahan, A. Snyder, P. Chi, D. Danila, M. Gounder, J. J. Harding, M. D. Hellmann, G. Iyer, Y. Janjigian, T. Kaley, D. A. Levine, M. Lowery, A. Omuro,

M. A. Postow, D. Rathkopf, A. N. Shoushtari, N. Shukla, M. Voss, E. Paraiso, A. Zehir, M. F. Berger, B. S. Taylor, L. B. Saltz, G. J. Riely, M. Ladanyi, D. M. Hyman, J. Baselga, P. Sabbatini, D. B. Solit, and N. Schultz. OncoKB: A Precision Oncology Knowledge Base. *JCO Precis Oncol*, 2017, Jul 2017.

[23] S. Chakraborty, A. Arora, C. B. Begg, and R. Shen. Using somatic variant richness to mine signals from rare variants in the cancer genome. *Nat Commun*, 10(1):5506, 12 2019.

**Figure SN1:** Oncotree is the tumor-type ontology encompassing all the possible tumor-type terms in our analysis (IntOGen). The outer nodes represent the most specific tumor-types in the hierarchy. Notice the central nodes for “CANCER”, and its children, “SOLID” and “NON-SOLID”, highlighted with different colors.

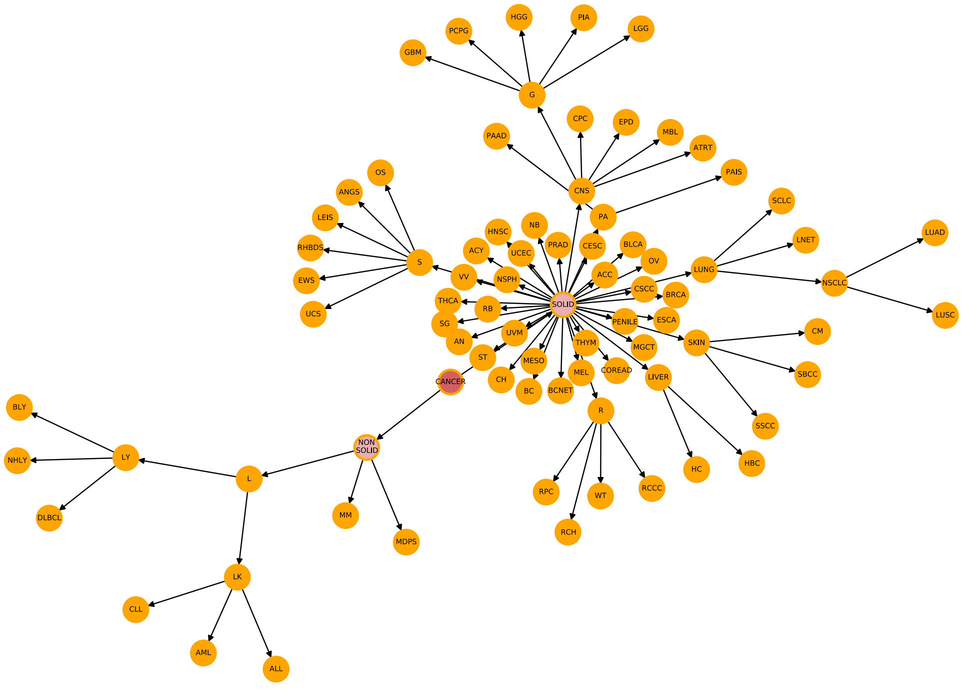

**Figure SN2:** The learning curves for Training (blue) and Test (red) show the log-loss (cross-entropy) for every training iteration for the 50 classifiers that make every model. Dashed vertical lines connect Train-Test pairs of points arising from the same classifier. Even though some classifiers attain the maximum number of estimators, the Test performance improvement in these cases is relatively weak. The panels show: **(a)** EGFR in lung adenocarcinoma (LUAD); **(b)** TP53 in colorectal cancer (COREAD); **(c)** PIK3CA in breast adenocarcinoma (BRCA); **(d)** CTNNB1 in hepatocellular carcinoma (HC).

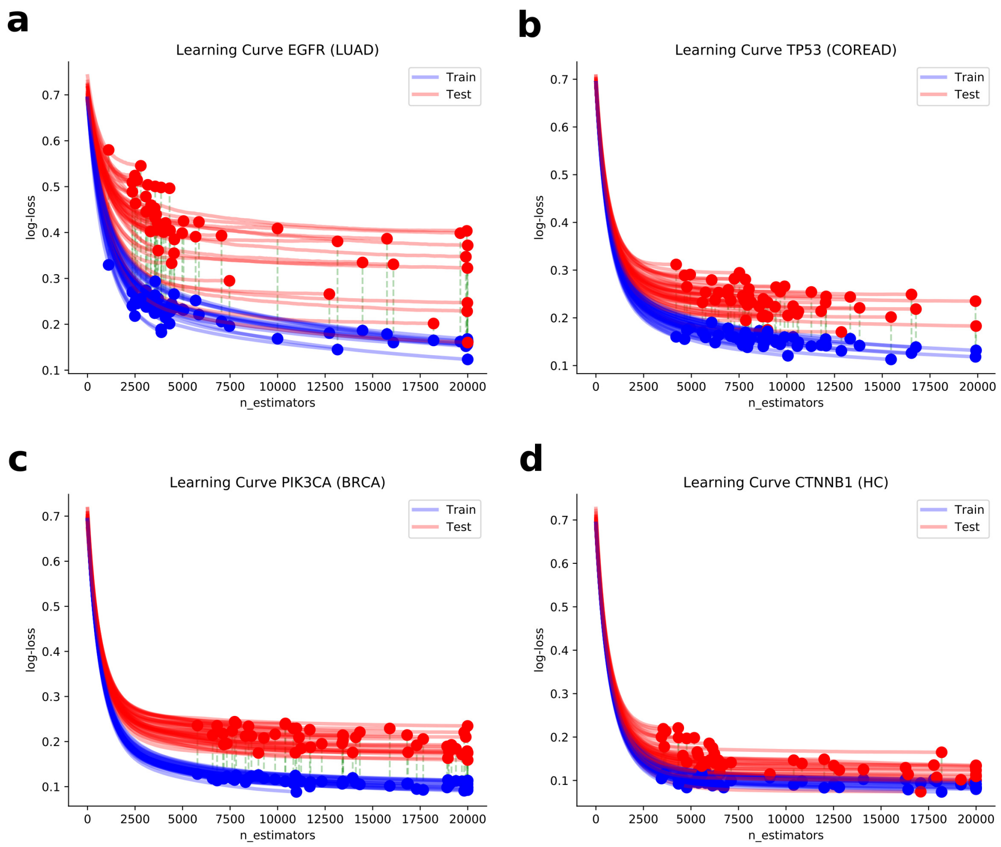

**Figure SN3:** **(a)** The `boostDM` model selection strategy is reminiscent to searching with the dials of a clock. After setting the gene value we scan through all the available tumor-type values upstream of the Oncotree. We move one step the gene value only if we restart the tumor-type value. **(b)** Although the model of FBXW7 in uterine carcinosarcoma (UCS) has mean 0.85 auROC, the small size of the training dataset, witnessed by a small size of the cross-validation Test sets, precludes a stable performance. **(c)** The quality criteria is first attained with the more general model for FBXW7 in the context of solid tumours (SOLID) where FBXW7 is a driver.

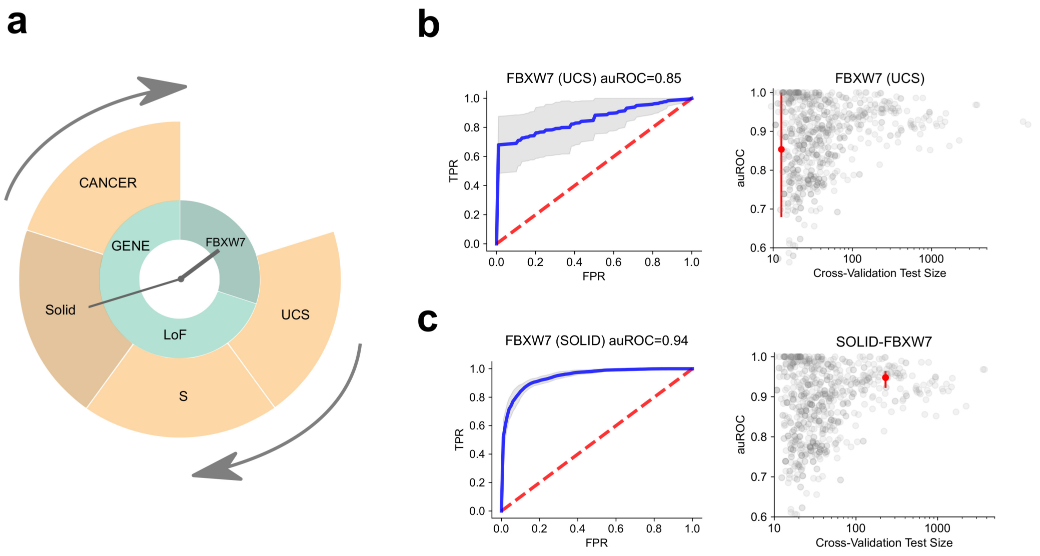

**Figure SN4:** **(a)** Fitness score for all missense mutations in PTEN (VEP.92 canonical transcript) reported in [18] divided in two groups depending on the `boostDM` forecast with the specific model for uterine corpus endometrial carcinoma. **(b)** ROC curve arising from (a) yields a modest 0.71 auROC. **(c)** auROC for all five tumor-types in which PTEN specific models are available.

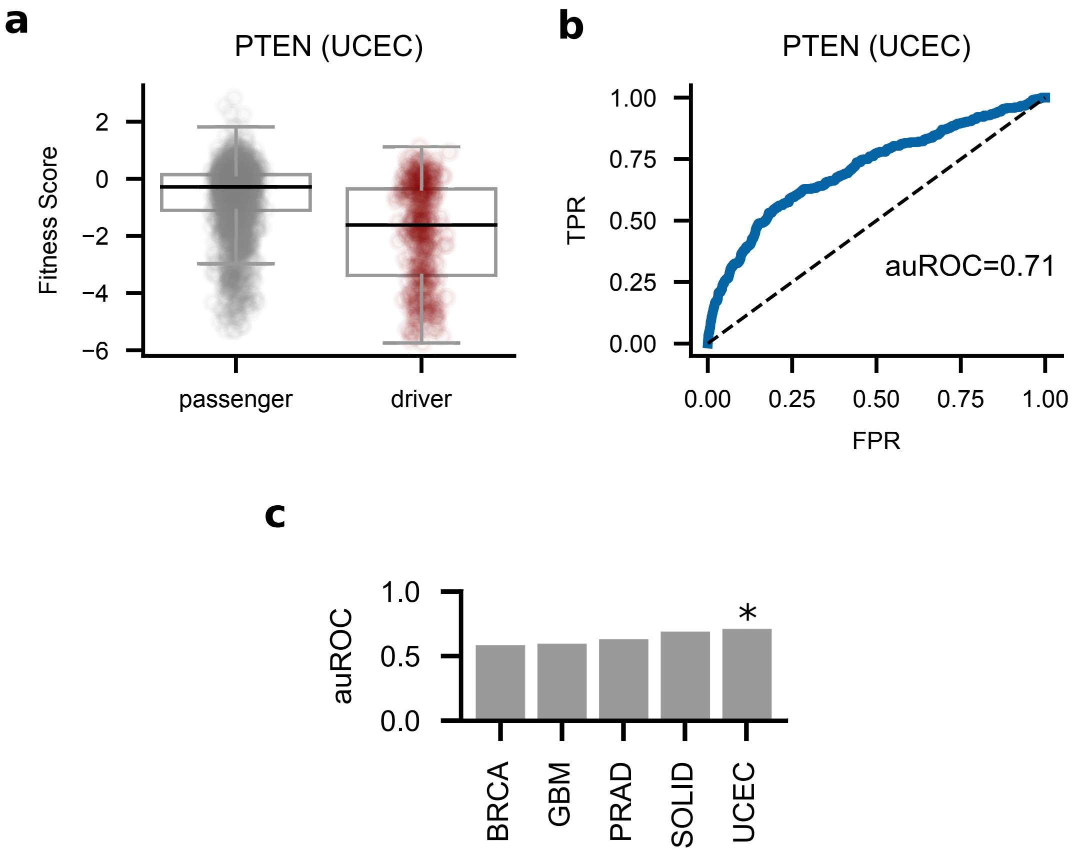

### Supplementary Tables

**Table S1. Description of datasets of somatic mutations of tumor types employed in the study.**

**Table S2. Oncotree relationships between tumor types**

**Table S3. Signatures included for signature fitting with deconstructSigs**

### Supplementary Data

**Supp Data 1. File containing the mutation probability of cancer gene-tumor type combinations**
